# Early warning signals for emerging infectious diseases

**DOI:** 10.1101/2025.03.03.641350

**Authors:** Qinghua Zhao, Nuno Moniz, Egbert H. van Nes, Marten Scheffer, Alexis Korotasz, Carly Barbera, Jason R. Rohr

## Abstract

Establishing early warning systems for infectious disease outbreaks could save millions of lives by enabling rapid response and containment. One promising approach draws on the concept of critical slowing down (CSD)—a phenomenon in which complex systems lose resilience before tipping points—detected using resilience indicators (RIs) derived from the statistical properties of time series. While disease outbreaks may exhibit such early warning signals of critical transitions, most prior applications of CSD theory to global health have been limited to single diseases or locations, without broad assessments of predictive accuracy or lead time—the interval between the detection of warning signals and the onset of an outbreak. To address these limitations, we integrate CSD theory with time-to-event analyses to evaluate the predictive performance of 17 RIs across 31 infectious diseases in 134 regions worldwide. We find that both RIs and time-to-event analyses provide ample time to implement control measures, reliably anticipating outbreaks with a mean lead time of 17–21 days. Lead time was greater for pathogens with longer incubation periods and in regions with higher Human Development Index. Additionally, temperature and precipitation exhibited unimodal effects on lead time predictions for vector-borne and viral diseases. These findings highlight the value of incorporating socio-environmental drivers into outbreak forecasting models and lay the foundation for a local-to-global early warning system capable of guiding proactive public health interventions.

## Introduction

Infectious diseases frequently re-emerge, driven by factors such as shifting human movement patterns, waning immunity, or reduced vaccine uptake (*1*). These resurgences cause millions of deaths worldwide each year; therefore, the ability to predict infectious disease outbreaks weeks in advance could save countless lives and significantly improve global quality of life (*2*). Several approaches can theoretically be used to anticipate outbreaks (*2–4*), such as applying dynamical mathematical models, monitoring Google searches and social media posts, wastewater surveillance for pathogens, and calculating generic, model-free statistical properties of time series as early warning signals (EWSs) of complex systems approaching critical transitions or “tipping points” (*5–7*). The value of using statistical properties of time series as EWSs is that they tend to require less effort and money than parameterizing dynamical models or monitoring wastewater for pathogens, and have regularly out-performed Google search and social media surveillance (*2*). Additionally, across many complex systems, statistical properties of time series are well linked to critical slowing down or reduced resilience (i.e., the ability to maintain normal stabilizing dynamics, e.g., a disease-free state, when subjected to disturbances) as systems approach critical transitions (*2*), which is why these types of EWSs are often referred to as resilience indicators (RIs). Measuring a system’s slowed recovery from a perturbation can be challenging, so statistical properties of time series, such as changes to the variance or autocorrelation, are used as RIs or proxies of critical slowing as systems approach tipping points (*2*).

Importantly, it is well documented that there were RIs in time series of past earthquakes, state shifts in ocean circulation systems and climate, and collapses of the global financial market, societies, and ecosystems (*5–7*). The application of RIs to human health has been more limited, but RIs have been detected for asthmatic attacks, mood depression, and epileptic seizures (*5–7*).

Recently, there has been interest in applying resilience theory to anticipate infectious disease outbreaks (*1*, *2*, *8–11*). Mathematically, the critical transition in models of infectious diseases is a piecewise continuous transcritical bifurcation, whereby self-limiting, stuttering chains of infection that inevitably go extinct shift to major outbreaks of sustained human-to-human transmission (*1*, *9*). When the reproduction number, R (the number of people in a population who can be infected by an individual at any specific time), is below the critical threshold of 1, the system is stabilized at a relatively disease-free state (i.e., few cases are observed), whereas above 1, the disease-free state becomes unstable and the introduction of a single infected individual can trigger an outbreak (*2*). Simple models of disease emergence show evidence of critical slowing down across a wide range of parameter values (*1*, *9*) and under varying conditions of ignorance and uncertainty (*12*, *13*), suggesting that RI-based EWSs for disease outbreaks might be common.

Recently, RIs were detected for malaria resurgence in Kenya (*10*) and COVID-19 waves (*14*), but the application of RIs to shifts from endemic disease-free to epidemic disease-emergence states have been predominantly theoretical and studies of real-world data have focused mostly on single diseases, regions, and RIs with inconsistent success (*1*, *2*, *8*, *11–13*). Hence, the use of RIs to offer reliable EWSs across infectious diseases remains equivocal. Additionally, little is known about the combinations of multiple RIs that could maximize EWS performance (Fig. 1), the performance of RIs across different types of diseases, or the traits of pathogens and locations that affect outbreak predictions and lead times (how early an outbreak is anticipated; Fig. 1), the latter of which is critical because it determines the amount of time communities have to prevent or mitigate an impending outbreak (but see (*9*)). Another major knowledge gap is that, unlike dynamical models or wastewater and social media monitoring, traditional RI-based approaches do not actually predict the probability or the timing of a critical transition because they never treat the timing of the actual transition (i.e., outbreak) as the dependent variable. Rather, RI-based approaches use a significant positive relationship between the RIs (dependent variable) and time before the critical transition (independent variable) as an indirect estimate of an impending state shift in the system, despite there being a well-established class of statistical analyses for studying how long it takes for a binary event to happen, which is referred to as time-to-event, failure time, or survival analyses (Fig 1).

**Figure 1.**
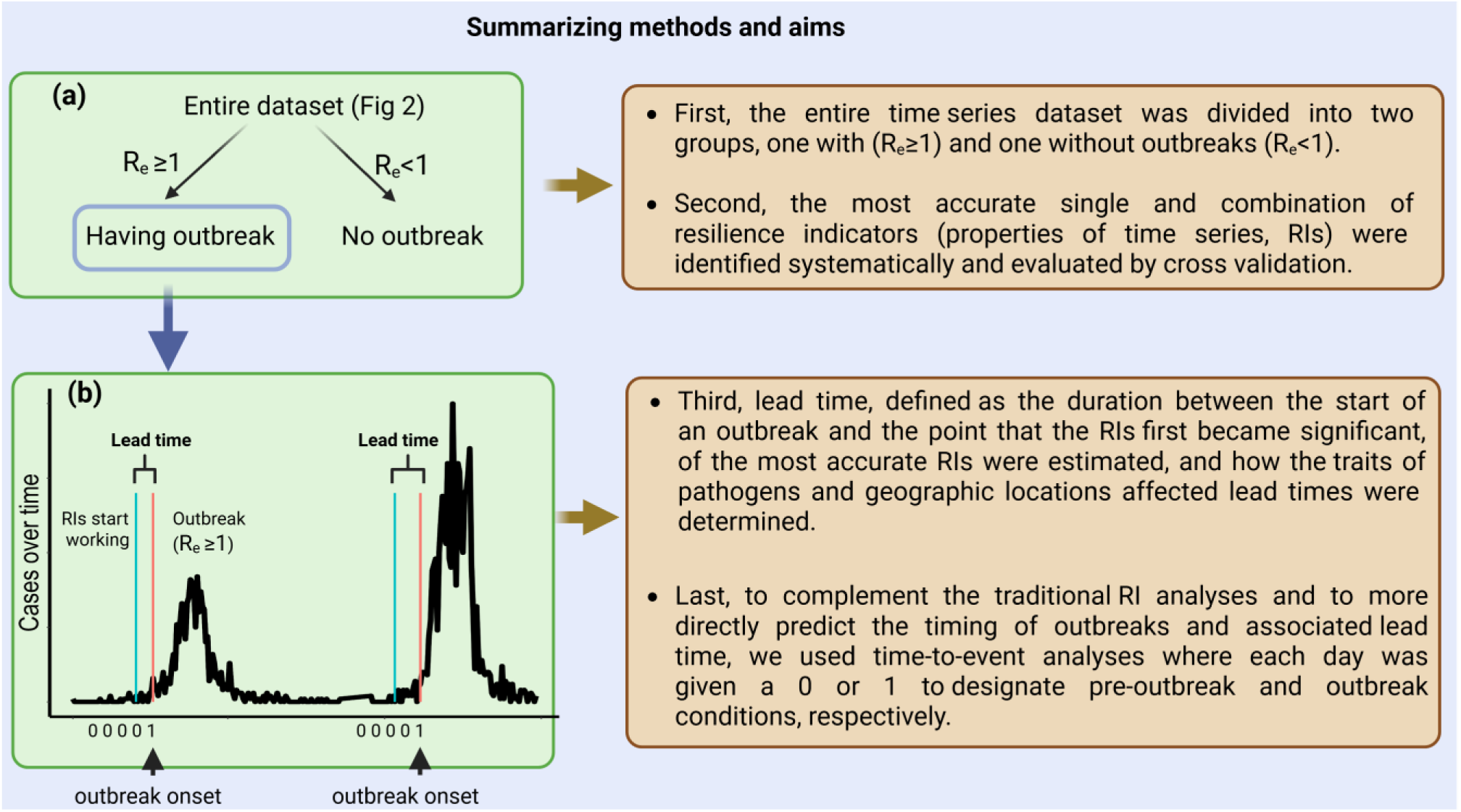
An overview of the data and methods used in this paper. **a.** The entire dataset of time series was divided into two groups, those with (i.e. the average number of secondary infections caused by a single infected individual at a given time point, 𝑅_𝑒_ ≥ 1; red line at **b**) and without outbreaks. **b**. Next, the most accurate resilience indicator (RI) was identified by considering single or combinations of RIs (see Methods section) with accuracy determined by taking the area under the receiver operating characteristic curve of 100 runs of a 5-fold cross validation. **b**. Once the most accurate RI was established, we quantified lead time or how early an outbreak could be anticipated (i.e., days between the start of an outbreak and the point that the RI first became significant). Next, we tested whether lead time was affected by the traits of the pathogens and geographic locations of the outbreak. Last, given that traditional RI analyses only indirectly predict outbreaks by testing correlations between RIs and time, time-to-event analyses were employed to more directly predict the timing of outbreaks and lead time.

To fill these gaps in understanding, we analyzed a high-resolution global dataset of daily incidence for 31 human infectious diseases across 134 regions worldwide (Fig 2), comprising 302 outbreaks (see Materials and Methods for a list of diseases). Among these 31 diseases, 64% were zoonotic, 58% were zoonotic diseases with a wildlife reservoir, 35% were vector-borne, and 54%, 35%, and 5% were caused by viruses, bacteria, and protozoa, respectively (Fig. 2). Using this dataset, we tested 17 RIs in total, 11 commonly used RIs, 4 RIs identified from machine learning and new modelling techniques, and 2 RIs specifically designed for time series influenced by seasonal forcing. To test for the significance of the RIs, we advanced a moving window forward towards the outbreak, testing for a significant positive association between the RIs and time in each window based on a Kendall 𝜏 rank correlation statistic (see Material and Methods). The performance of the RI in predicting outbreaks was evaluated based on its true positive and negative rates (i.e., using the area under the receiver operating characteristic curve (AUC)) and 5-fold cross validation. For outbreaks that could be predicted using an RI, we calculated the lead time as the days between the first instance that 𝜏 was significant and the start of the outbreak (Fig. 1). In contrast to this traditional RI approach, we also used a time-to-event, Cox proportional hazards analysis to more directly predict lead time and test predictors of the timing of outbreaks.

**Figure 2.**
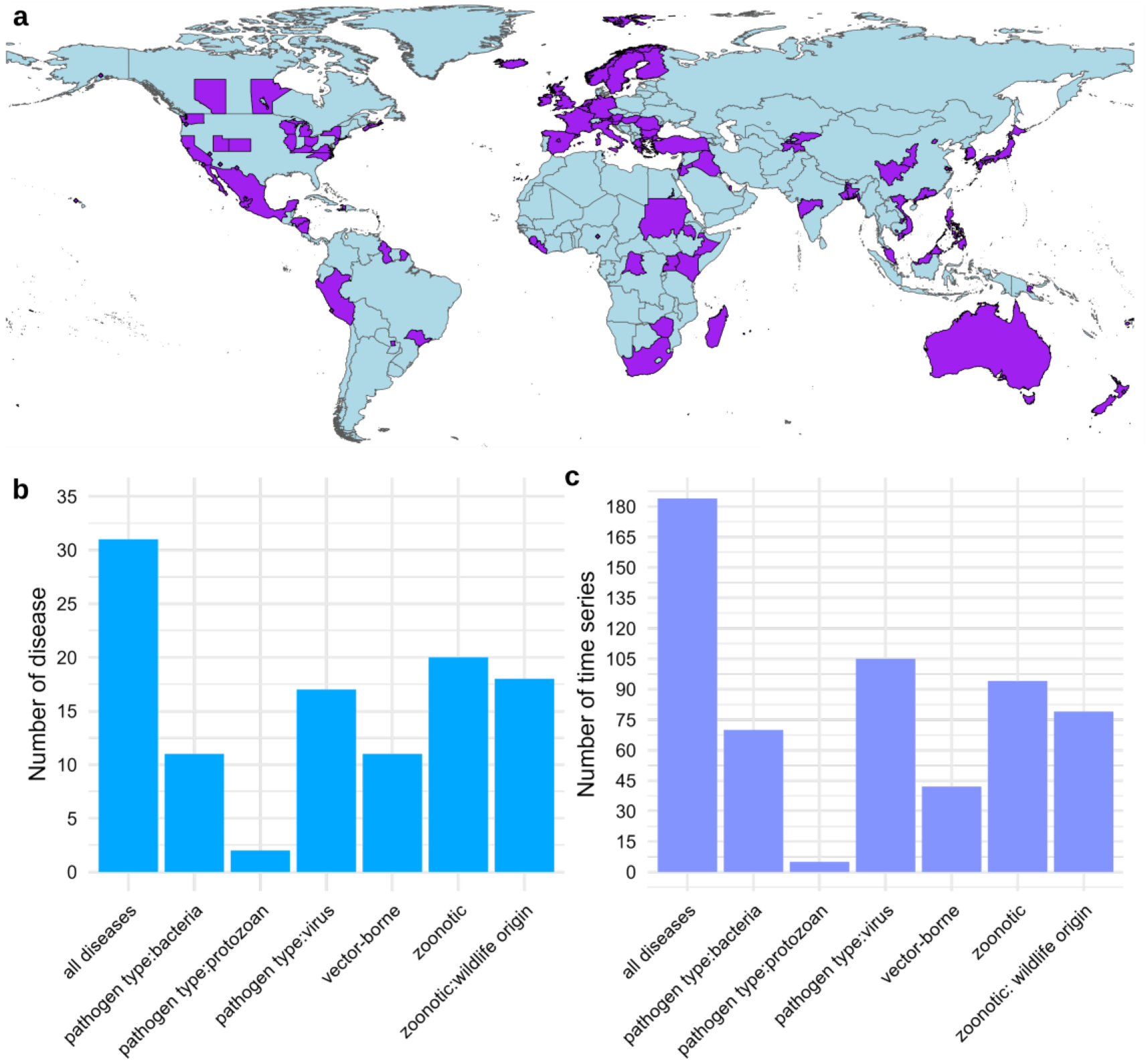
The studied locations (a), total number of diseases (b), and total number of time series (c) grouped by pathogen- and disease-type categories. See Table S4 for details of diseases, regions, and pathogen types.

## Evaluating resilience indicators and lead times

Across all 31 diseases, we found that the best performing single RI across moving window lengths was Standard Deviation (mean AUC=0.93 from cross validation, window length=10 days) (Fig 3a-b, S1). Importantly, the performance of Standard Deviation extends beyond what can be explained simply by trends in the mean incidence. Indeed, trends in the mean incidence exhibit lower maximum predictive power (AUC=0.70, Fig S2) compared to those of Standard Deviation or the first-order autoregressive model (AR-1; AUC=0.87, Fig S1h). While the Standard Deviation provides the highest overall predictive performance, the observed increase in AR-1 prior to outbreaks is particularly notable, as autocorrelation is widely regarded as a robust indicator of CSD (*15*).

**Figure 3.**
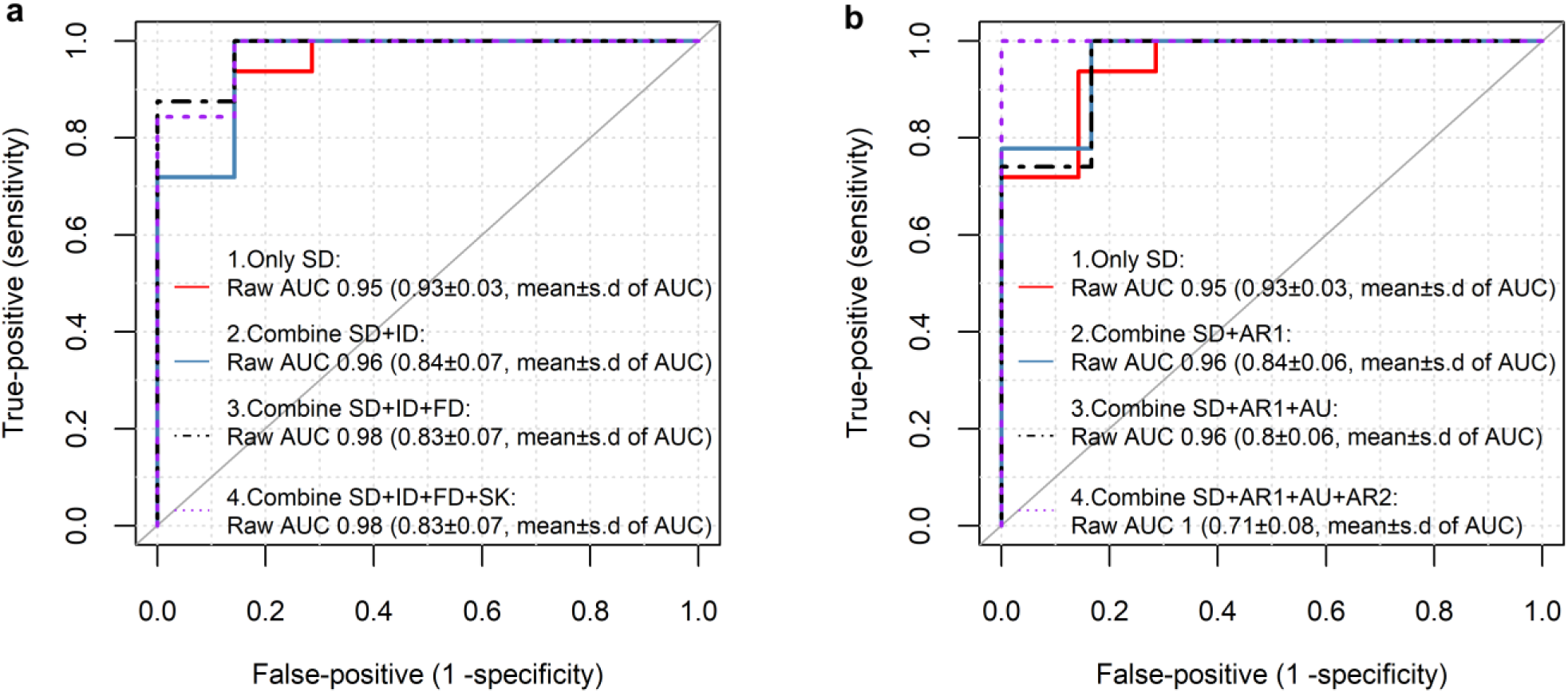
The accuracy of resilience indicators (RIs) as early warning signals (EWSs) of outbreaks across 31 diseases, examined using mean area under the receiver operating characteristic curves (AUC) and cross validation. All RIs used the window length (i.e. 10 days) that maximized the AUC for the most accurate RI (i.e. Standard Deviation; mean AUC=0.93 from cross-validation) (**a-b**) across all RIs and window lengths (Fig S1). We sequentially added either the next best performing single RI (**a**) or the next most orthogonal RI (i.e., least correlated based on Principal Component Analysis) to the best performing RI (**b**) until we tested up to four total RIs in our models. Importantly, sequentially adding the next most orthogonal RI to the model ensures that the baseline AUC goes up with each RI addition, but it does not guarantee that the mean AUC from the cross validation will continually rise with each added RI. Standard deviation: SD, index of dispersion: ID, first differenced variance: FD, skewness: SK, autocorrelation at lag 1: AR1, autocovariance: AU, autocorrelation at lag 2: AR2. The AUC values in parenthesis at **a-b** represent the mean and s.d from cross-validation.

Therefore, the concurrent rise in both Standard Deviation and AR-1 suggests that the predictive power of our early warning approach is indeed rooted in the generic phenomenon of CSD. Combining the Standard Deviation with other resilience indicators (RIs) did not enhance predictive accuracy, as the mean AUC from cross-validation did not improve when Standard Deviation was combined with either the top-performing RIs or those least correlated with it, despite improvements observed in the raw (non-cross- validated) AUC (Fig 3a, 3b). Hence, using the raw AUC rather than the mean AUC from cross validation can result in overfitting and thus overly optimistic estimates of model performance. For the best RI (Standard Deviation), the mean lead time was 17.7 days (±1.3 s.e.; Fig. 4) across times series and diseases. The time-to-event analyses, which captures both the likelihood of an outbreak and its timing, also produced a dependable model, accounting for 65.6% (marginal *R*^2^) of the variation in the hazard of an outbreak (Fig. S3). The higher 𝜏 value from the best RI (Standard Deviation) was negatively associated with the time to an outbreak in the time-to-event analyses (Fig. S3b) and thus a higher hazard of an outbreak (Fig. S3a), reflecting the consistency between the traditional RI and time-to-event models. The time-to-event analysis estimated a mean lead time of 21.3 days (±1.2 s.e.; Fig. S4), which is 3.6 days longer than the estimate provided by the traditional RI approach. Importantly, given that major outbreaks do not happen immediately after *R*_e_ = 1 (*9*), the actual lead time likely exceeds these estimates. Overall, these results suggest that both RI and time-to-event analyses reliably warn of upcoming outbreaks, and the average warning provided by RIs of 17-21 days or greater is enough time to mitigate, and possibly even prevent, outbreaks.

**Figure 4.**
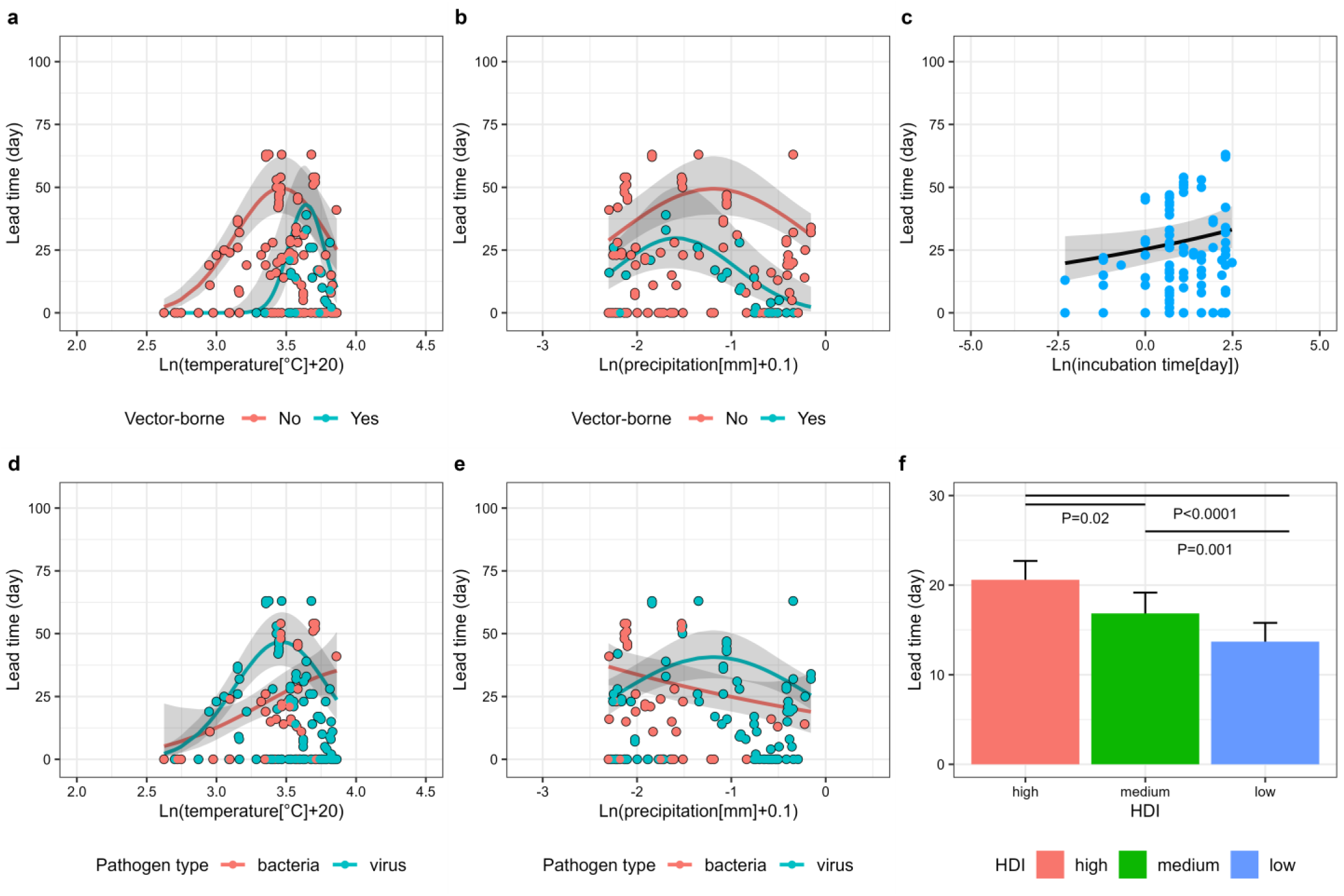
A display of the main effects and interactions that were significantly associated with lead time. The bold lines and gray bands depict best-fit lines and 95% confidence bands (or standard error in panel f), respectively, derived from the predicted values of the final statistical model that is available in Table S1. In panel **f**, the significance between groups was assessed using the emmeans function from the emmeans package. For the effects of time series length and population density on lead time, refer to Fig S7-S9. Statistical results see table S1. HDI represents the Human development index.

Given that many datasets provide weekly rather than daily case report data, we reconducted our models averaging the daily data of each week. Notably, the coarser- scale weekly data resulted in lower AUC values (Fig. S5) and a mean lead time of 50.6 days (Fig. S6), which is > 30 days longer than the lead time from the daily data. This indicates that temporally coarse data resolution can provide an overly optimistic estimation of lead times for outbreak mitigation and prevention. Therefore, the use of finer-scale daily data is highly recommended for more reliable EWS assessments and lead time estimates.

## How traits of pathogens and outbreak locations affect both lead time and outbreak risk

Next, using the daily data, we sought to identify the traits of pathogens and outbreak locations that could explain variation in the time to outbreak occurrence and lead times predicted by the best RI (Standard Deviation). We hypothesized that outbreaks caused by pathogens with shorter incubation times (the interval between exposure to a pathogen and the appearance of the first symptoms) would be harder to anticipate and would have shorter lead times than those with longer incubation times. Temperature and precipitation were expected to exhibit hump-shaped relationships with time to outbreak occurrence and lead time, as these environmental factors are known for unimodal effects on transmission dynamics and pathogenesis (*16*), particularly for vector-borne pathogens, all of which have ectothermic hosts that are often more sensitive to climatic factors than endothermic hosts. Additionally, given that vector-borne pathogens tend to have greater frequency- than density-dependent transmission (*17*), we postulated no relationship between lead time and human population density for these pathogens. Finally, given that more developed areas have greater outbreak surveillance, health infrastructure, and case reporting, we hypothesized that regions with a higher Human Development Index (HDI; a statistical composite of life expectancy, years of education, and per capita income) (*18*) would have a longer lead times than regions with lower HDIs. Additionally, we predicted that HDI would be negatively associated with the risk of disease outbreaks for vector-borne diseases because they are often effectively combatted by costly insecticide applications and homes with windows or screens (*19*, *20*). To test these *a priori* hypotheses, we developed a baseline generalized linear mixed effects model for lead times predicted by the best RI (Standard Deviation) and a Cox proportional hazards model for time to outbreak occurrence, and removed nonsignificant factors through a backwards selection process to reach a final model (see Materials and Methods, Table S1-S2).

These analyses revealed considerable variation in lead times and time to outbreak occurrence across the time series and 31 diseases. For example, as predicted, pathogens that spread rapidly because they have short incubation times also had short lead times, with each natural log increase in days of incubation time increasing lead time by 0.11 ±0.05 day (mean±s.e) (Fig. 4c). Additionally, on average, vector-borne and viral pathogens offered shorter lead times (12.5 and 16.1 days) than non-vector-borne and bacterial pathogens (19.2 and 19.0), respectively (Fig. 4a, 4b). However, when temperature and precipitation were near their mean values, lead times peaked for both vector-borne and non-vector-borne pathogens, and vector-borne pathogens had shorter lead times than non-vector-borne pathogens (Fig. 4a, 4b). In fact, at mean temperature and precipitation, lead times were 40.0 and 27.0 days, respectively, for vector-borne diseases, and approximately 50.0 and 50.0 days, respectively, for non-vector-borne diseases (Fig. 4a, 4b). Additionally, vector-borne diseases occurred under more narrow temperature conditions and peaked at warmer temperatures (20.4 °C vs 13.3 °C) than non-vector-borne diseases (Fig. 4a), consistent with the well-documented sensitivity of vector-borne pathogens to environmental conditions and greater frequency of vector-borne diseases in warmer than cooler regions (*16*).

Whereas lead times for both vector- and non-vector-borne diseases responded unimodally to temperature, only viral pathogens were unimodally associated with temperature, peaking near observed mean temperatures (13.1 °C) (Fig 4d). In contrast, bacterial pathogens were associated positively with temperature for both lead time and time to outbreak occurrence (Fig. 4d, 5b). Consistent with these findings, several studies documented hump-shaped relationships between arboviruses and temperature (*16*), whereas bacterial biological rates (*21*) and infections in humans regularly show positive associations with environmental temperature (*22*). Precipitation exhibited a unimodal relationship with both lead time and time to outbreak occurrence for vector- borne and non-vector-borne diseases (Fig. 4b, 5c) and a unimodal relationship with lead time for viral pathogens (Fig. 4e), consistent with previous studies documenting unimodal relationships between precipitation and these pathogen types (*23*, *24*). In contrast, precipitation was negatively associated with lead time for bacterial infections (Fig. 4e), aligning with previous research documenting that higher precipitation levels facilitate more rapid transmission of bacterial pathogens (*22*, *25*).

As predicted, lead time was associated positively with HDI, presumably because wealthier locations invest more in health infrastructure and pathogen surveillance that can facilitate earlier detection of human cases of disease (Fig. 4f). This positive association between lead time and HDI was significant across all diseases (Fig. 4f). The time-to-event analysis further revealed an HDI-by-vector-borne-disease interaction, with vector-borne diseases showing a positive association between HDI and time to outbreak occurrence (Fig. 5d). In fact, a shift from the low to high HDI category more than halved the hazard of a vector-borne disease outbreak (Fig. 5d). These results are consistent with vector-borne diseases being reduced by human development more than non- vector-borne diseases, thus lengthening the time to vector-borne disease outbreaks. For example, many vector-borne diseases can be effectively managed by costly home improvements and insecticide applications (*19*, *20*). Hence, development can enable earlier detection and thus mitigation and prevention of environmentally mediated diseases.

**Figure 5.**
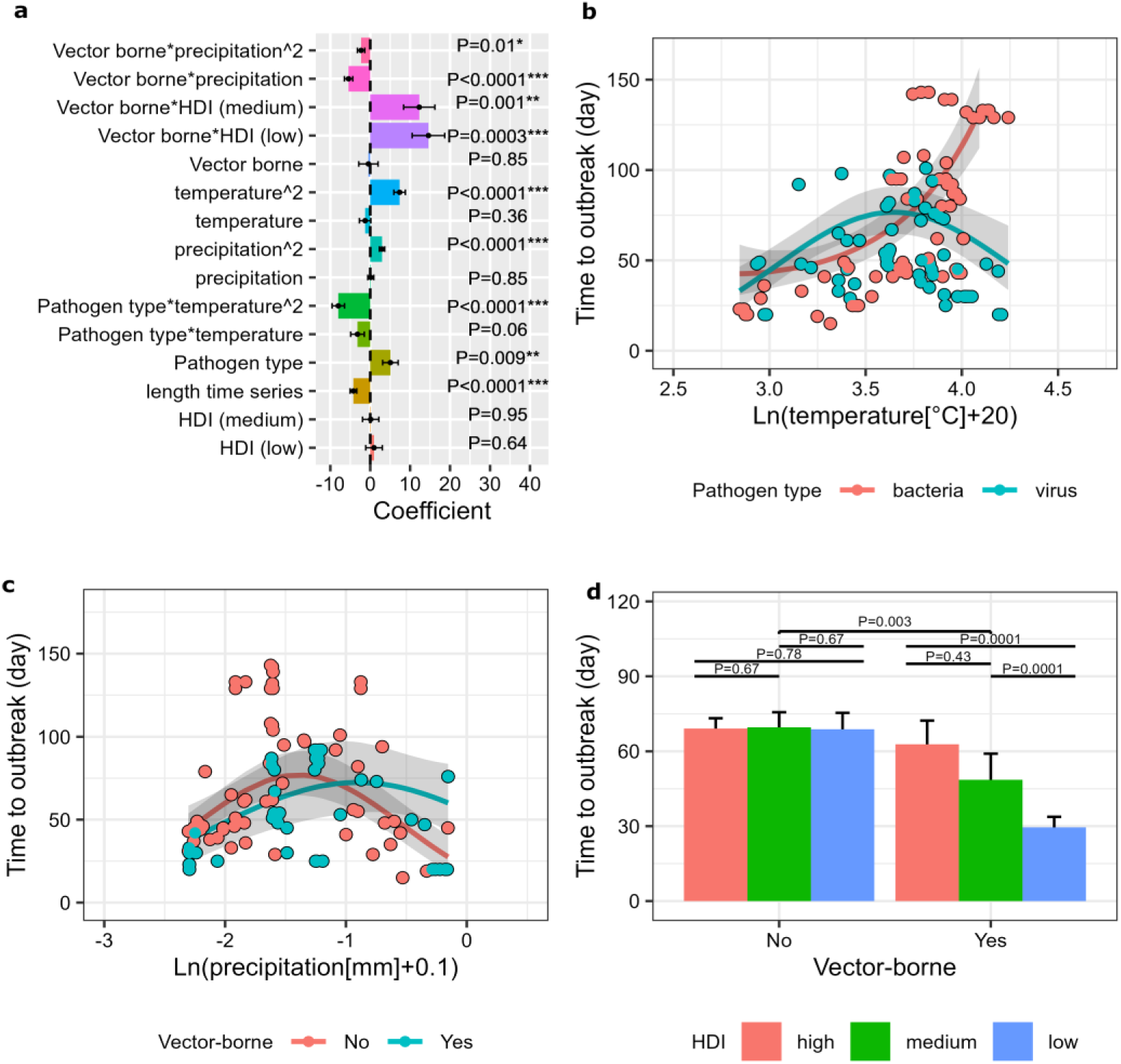
Coefficients ± s.e (a) and plots of significant predictors (b-d) from a Cox proportional hazards mixed model examining the effects of various factors on time to outbreak occurrence. **a**, A positive coefficient signifies an increased hazard of outbreak occurrence, and consequently a reduced time until an outbreak. A significant quadratic coefficient indicates a second-order polynomial function between a predictor and hazard of outbreak occurrence. The bold lines and gray bands depict best-fit lines and 95% confidence bands (or standard error in panel f), respectively, derived from the predicted values of the final statistical model that is available in Table S3. **d**, the significance between groups was assessed using the emmeans function from the emmeans package. Significant effects are indicated by *P <0.05, **P < 0.01, ***P < 0.001. HDI represents the Human development index.

Time series length was associated positively with lead time (Fig. S7a) and time to outbreak occurrence (Fig. S7b), suggesting that early initiation of monitoring and case reporting (*26*, *27*), the availability of longer historical datasets (*28*), or an enhanced commitment to outbreak surveillance (*29*) might enable stakeholders to extend lead times before an outbreak. Additionally, neither outbreak type (first or subsequent in the time series) nor human population density had significant effects on lead time or the hazard of outbreak occurrence (Fig. S8 & S9). However, we admonish against assuming that these factors do not affect these endpoints, as increased statistical power or a different subset of human diseases could produce different results.

While our study provides critical advancements in the development and application of resilience theory for predicting disease outbreaks, we acknowledge potential misclassifications of case data stemming from reporting biases, diagnostic inaccuracies, or surveillance inconsistencies, which may introduce uncertainties and weaken the robustness of warning indicators. Importantly, these misclassifications should only amplify random errors in our analyses that would reduce the likelihood of detecting RIs that warn of impending outbreaks. Nevertheless, we discovered EWSs despite this impediment. Another important limitation of our analyses is that they are restricted to gradual endemic states of pathogen transmission rather than abrupt emergence of outbreaks involving new strains (e.g., a reassortant influenza virus) or pathogens (e.g., de novo spillover of a human-to-human transmissible pathogen from a zoonotic reservoir) to a region (*11*). Expanding the framework to incorporate RIs for new pathogens, such as those that spillover from wildlife to humans, and artificial intelligence (*11*, *30*) would enhance its utility in broader public health contexts.

## Implications and Conclusions

Our study provides a significant advancement in the development and application of EWSs for infectious disease outbreaks. By systematically evaluating multiple RIs across a diverse set of diseases and geographic locations, we identified key predictive factors that enhance outbreak forecasting. Shifting to time-to-event analysis from the traditional RI-based approach improved both the accuracy of outbreak predictions and lengthened lead time, providing a more reliable framework for early intervention.

Importantly, we demonstrate that pathogen characteristics, climate, and socio- economic factors significantly influence lead time and outbreak timing, highlighting the need for context-specific forecasting models. Our findings show that pathogen incubation time is associated positively with lead time. Additionally, higher HDI regions experience longer lead times and reduced outbreak risks for vector-borne diseases, suggesting that public health infrastructure and surveillance capacity play crucial roles in disease prevention. Furthermore, we emphasize the need for high-resolution temporal data, as coarse-scale weekly data may lead to overly optimistic lead time estimates, potentially compromising timely responses.

This research lays the foundation for a global disease forecasting system that integrates real-time data streams and model-free statistical techniques to enhance predictive accuracy (*11*). Future work should focus on expanding the range of predictive indicators and developing interactive platforms for real-time outbreak monitoring. By improving our ability to anticipate and mitigate outbreaks, our study contributes to strengthening public health preparedness and minimizing the societal and economic impacts of infectious diseases.

## Methods

### Disease dataset

For our analyses, we used the subset of the Analytics for Investigation of Disease Outbreaks (AIDO) dataset (https://aido.bsvgateway.org/) with daily reported disease case data and time series ≥1 week. This resulted in 31 diseases across 134 regions (i.e. either country or administrative level e.g. municipality and province; Fig 2), comprising 184 time series (Fig 2) with an arithmetic mean length of 117 days (±14 s.e.) (see Table S4 for the corresponding regions and time series length for each disease). The 31 diseases included in this dataset are Anthrax, Brucellosis, Campylobacteriosis, Chikungunya, Cholera, Dengue, Ebola, Gastroenteritis, Lassa Fever, Leptospirosis, Malaria, Marburg, Measles, Middle East Respiratory Syndrome, Monkeypox, Nipah, Norovirus, Novel Influenza A, Pertussis, Plague, Polio, Q Fever, Rift Valley Fever, Rubella, Salmonellosis, Shiga Toxin-Producing E. Coli, Shigellosis, Tularemia, West Nile Virus, Yellow Fever, Zika.

### Socio-economic and eco-environmental factors

We gathered covariates, all of which were matched to each time series and corresponding location in the disease dataset. Daily temperature (°C) and precipitation (mm) were retrieved from NOAA’s National Centers for Environmental Information (*31*). Annual human development index (HDI) for each 5 arc-minute gridded unit (latitude and longitude) was sourced from Kummu et al. (*18*). and subsequently aggregated to the respective country or administrative level using arithmetic means. Yearly population density (people per km^2^) was retrieved from the Center for International Earth Science Information Network database (*32*). Across all studied locations, the yearly resolution was used for HDI and population density because this was the only temporal resolution available. For each pathogen at a given location, the incubation time (day) was retrieved from the literature. Population density and incubation time were natural log transformed to ensure normality. Precipitation data were transformed using ln(x + 0.1) to accommodate zero values, while temperature data were transformed using ln(x + 20) to adjust for temperatures below −10°C in certain regions (e.g. Nordic countries such as Finland).

### Traditional RI analyses

Using the methods of Cori et al. (*33*) (because it outperforms others approaches (*34*); for details of these methods, see Supplementary Information 1), we first delineated the start of each outbreak in each time series as the point when 𝑅_𝑒_(i.e. the average number of secondary infections caused by a single infected individual at a given time point) (*11*, *33*, *34*) was ≥1. Thus, each time series was divided into no outbreak and outbreak sections (all code archived on GitHub (*35*)). Of the 184 time series, 27 did not have any days where 𝑅_𝑒_ was ≥1, representing no outbreak time series. Of the remaining 157 time series, 68 (43.3%) had a single outbreak, and 89 (56.7%) had multiple outbreaks, resulting in 302 outbreaks in total (see Fig S10 for the frequency of outbreaks). For each time series, we then utilized 11 commonly used RIs (standard deviation, skewness, kurtosis, coefficient of variation, first differenced variance, autocovariance, index of dispersion, density ratio, autocorrelation at lag 1, lag 2 and lag 3), 2 RIs specifically designed for time series influenced by seasonal forcing (wavelet filtering, wavelet reddening), and 4 RIs developed by machine learning, empirical dynamic modeling, or previously untested in the context of infectious diseases (relative dispersion, time series acceleration, the absolute value of dominant eigenvalue, and hurst exponent), resulting in a total of 17 RIs (*35*) (Table S5 and Supplementary Information 2).

Following Dakos et al. (*36*), we adopted a moving window approach to test for a positive association between each RI and time as it approached 𝑅_𝑒_ ≥ 1. Before conducting the moving window approach, each disease time series was detrended using Gaussian smoothing (*36*). The length of the moving window varied from 7 to 21 days (1 to 3 weeks) with increments of 1 day (Fig S1). The magnitude of the association between each RI and time in each window was measured using the nonparametric Kendall 𝜏 rank correlation statistic (*36*). The significance of the 𝜏 was determined by estimating the fraction of 100 null surrogates that performed better than the observed dataset using the method of Ebisuzaki (*37*). These randomized surrogate datasets have the same power spectra and autocorrelation structure of the original data but with randomized phases to eliminate spurious results because of the correlation in time.

Next, we employed receiver operating characteristic (ROC) curves to quantify the performance of each RI using 100 runs of 5-fold cross validation (i.e., 80% training data and 20% testing data), performed with the caret package (*38*). The cross-validation aimed to test out-of-sample model performance to determine if our models were underfit, overfit, or “well-generalized”. Cross-validation was performed to evaluate the predictive performance of RIs, rather than assess the fit to the raw case time series itself. For each of 100 runs, the area under the ROC curve (AUC) was computed to quantify the model’s discrimination ability, with a larger AUC indicating higher true positive and negative rates and thus higher accuracy. The arithmetic mean AUC obtained from 100 runs of cross-validation was taken as the final performance metric to assess the accuracy performance of each RI. Given that the length of the moving window can impact the AUC of an RI, the optimal moving window length was chosen for each RI by selecting the window length between 7 and 21 days with the highest mean AUC in the cross validations (Fig S1).

Once the most accurate single RI was determined, we tested whether combining up to four RIs would improve model predictions based on AUC and cross validation. Specifically, we forced all RIs to use the window length that maximized the AUC for the most accurate RI (i.e., Standard deviation with window length=10 days in Fig S1). Next, we conducted Principal Component Analyses (PCA) to determine associations among the RIs within this window length. Then, we sequentially added either the next best performing single RI or the next most orthogonal RI (i.e., least correlated based on PCA) to the best performing RI until we tested up to four total RIs in our models. Here, the multiple RIs were input as independent predictors within the train function of the caret package to implement cross-validation, thus taking a multiple regression approach to combining RIs. Importantly, sequentially adding the next most orthogonal RI to the model ensures that the baseline AUC goes up with each RI addition, but it does not guarantee that the mean AUC from the cross validation will continually rise with each added RI (Fig 3).

Once the most accurate single or combination of RIs across all 31 diseases was identified, we computed and analyzed the lead time (Fig. 1). The lead time was calculated as the days between the start of an outbreak (𝑅_𝑒_ ≥ 1) and the point that the RI or combination of RIs first became significant. Next, we employed a Generalized Linear Mixed Model (glmmTMB function in glmmTMB package (*39*)) with a negative binomial error distribution to test various *a prioiri* hypotheses regarding lead time highlighted in the main text. We tested the main effects of temperature, precipitation, HDI, pathogen incubation time, human population density, time series length, two pathogen traits (vector-borne or not; virus vs. bacteria), outbreak type (first outbreak in a time series or not), and all pairwise interactions between the pathogen traits, temperature, precipitation, HDI and human population density (Table S1). The parasite taxa were limited to viruses and bacteria, because our dataset had no fungi and helminths and because the sample size for protozoa was too small to justify analyses (Table S4). We compared the effects of first versus later outbreaks within time series to account for potential differences in lead time between initial and subsequent outbreaks. See the main text for the *a prioiri* hypotheses used to justify the interactions in this model. Additionally, we fitted the temperature and precipitation as quadratic polynomial functions, because most organisms have an optimal temperature and moisture (*16*), and treated disease types nested within regions as random intercepts. Additionally, HDI was classified into low (<0.55), medium (0.55–0.699), and high (≥0.7) categories, following the recommendation of The United Nations Development Programme (*40*). Figures were generated from the statistical model using the ggpredict function in the ggeffects package (*41*).

### Time-to-event analyses

To complement the indirect approach used by traditional RI analyses, we employed a mixed effects Cox proportional hazards time-to-event analysis (coxme function in coxme package (*42*)) to more directly test predictions of the timing of outbreak occurrence. We used the same model structure as described for the lead time analyses (Table S1). In these analyses, the response variable was a time series of 0 values indicating non-outbreak days (𝑅_𝑒_ < 1) and a single 1 value indicating the outbreak point where the outbreak began (𝑅_𝑒_ ≥ 1). Given that there is no reliable way to graph the time-to-event analyses incorporating random intercepts (*43–45*), we replicated the final Cox proportional hazards model in the glmmTMB package with negative binomial errors, verified that the direction of the effects and significance matched between the two models (final results see Table S2, S3), and then generated figures from the glmmTMB model (*39*) using the ggpredict function in the ggeffects package (*41*).

To directly estimate lead time rather than relying on indirect estimation as done by traditional RIs, we employed the parametric survival regression model (*survreg* function in survival Package) (*46*), i.e., a time to an event model. In the model, the predictor was the tau value (𝜏) of the best RIs (Standard Deviation), window length=10 days; Fig 3a-b). The response variable was a time series of 0 values indicating non-outbreak days (𝑅_𝑒_ < 1) and a single 1 value indicating the hypothetical time point at which the outbreak could be anticipated in advance. The lead time was computed as the days between the first instance that survival regression model was significant and the start of the outbreak. All analyses were conducted by R version 4.1.2 (*47*).

## Acknowledgements

We thank Prof. Stephen Carpenter for feedback on this manuscript. We thank the members of Rohr Laboratory of Ecology and Public Health at University of Notre Dame for discussion.

## Author Contributions

J.R.R. and Q.H.Z conceptualized the study. J.R.R. supervised the study. E.N.N and M.R. supported EWS analysis. N.M. helped with the selection of the RIs. Q.H.Z. performed the study and analyzed the data. J.R.R. and Q.H.Z. wrote the original draft. All the authors reviewed and edited the manuscript.

## Data and materials availability

All raw daily disease data used in this study are publicly available at ADIO (https://aido.bsvgateway.org/). All data, code, and materials used in the analysis are available on GitHub (https://github.com/QZhao16/EWs).

## Supplementary materials

### Supplementary Figures

**Figure S1.**
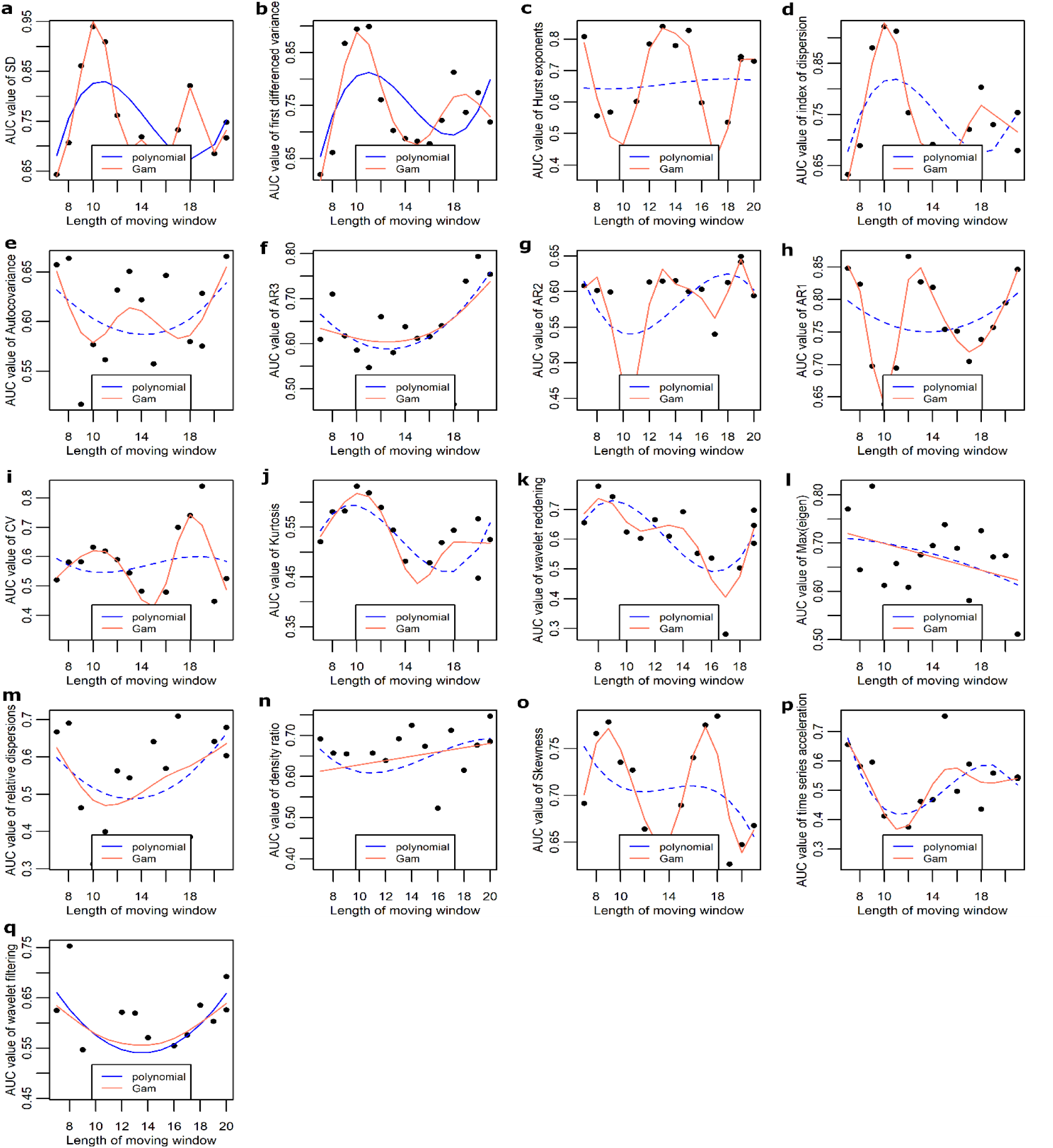
Across all 31 diseases, RIs accuracy (mean AUC from 100 runs of 5-fold cross validation) varies as a function of window lengths of 7 to 21 days. A higher AUC value (area under the curve) indicates a higher ability of a binary classifier to properly classify true positive and negative outcomes, thus representing higher accuracy for RIs. The red line represents the fit from a generalized additive model (gam function from mgcv package of R). The blue line represents a linear or quadratic fit from a linear regression model (lm function from the base package of R). The solid lines indicate significant associations, whereas dashed lines indicate non-significant associations.

**Figure S2.**
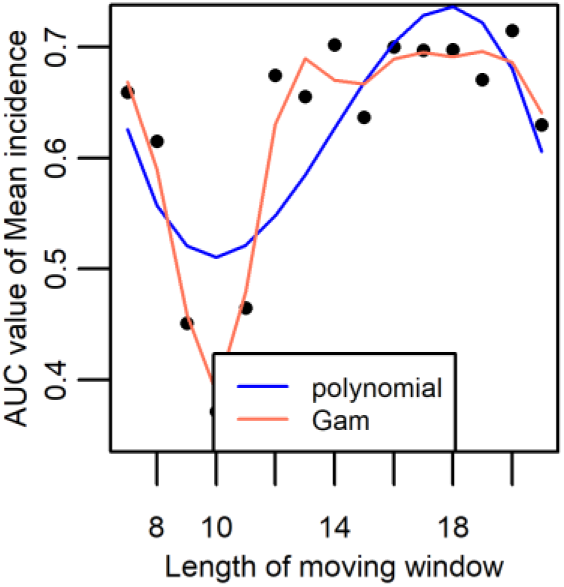
Across all 31 diseases, the accuracy (the AUC from 100 runs of 5-fold cross validation) of mean incidence varies as a function of window lengths of 7 to 21 days. All methods for calculating the area under the curve (AUC) for the mean incidence indicator are identical to those used for the 17 resilience indicators (see Methods and Fig S1). A higher AUC value (area under the curve) indicates a higher ability of a binary classifier to properly classify true positive and negative outcomes, thus representing higher accuracy for RIs. The red line represents the fit from a generalized additive model (gam function from mgcv package of R). The blue line represents a linear or quadratic fit from a linear regression model (lm function from the base package of R). The solid lines indicate significant associations, whereas dashed lines indicate non-significant associations.

**Figure S3.**
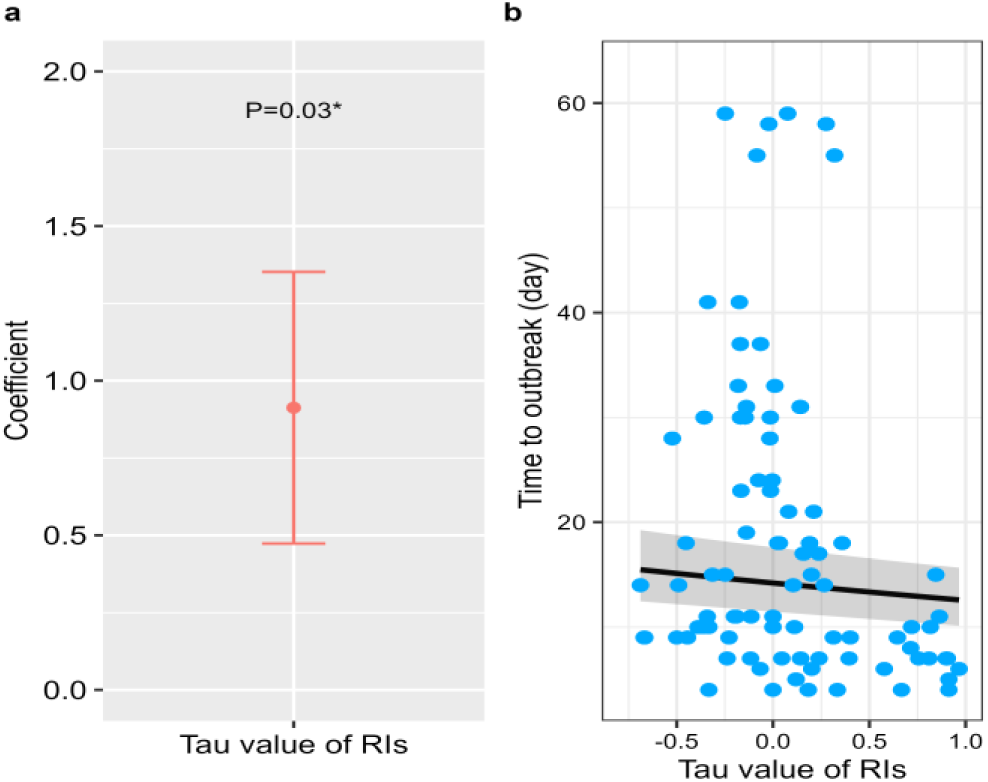
Coefficients ± s.e (a) and plots of tau value (𝝉) (b) from a Cox proportional hazards mixed model examining the effects of tau value (𝝉) from the best of RIs (Standard Deviation) on time to outbreak occurrence. In the model, we treated regions nested within disease types as random intercepts. The predictor was the tau value (𝜏). The response variable was a time series of 0 values indicating non- outbreak days (𝑅_𝑒_ < 1) and a single 1 value indicating the outbreak point where the outbreak began (𝑅_𝑒_ ≥ 1). **a**, A positive coefficient signifies an increased hazard of outbreak occurrence, and consequently a reduced time until an outbreak at (b). **b**, the bold lines and gray bands depict best-fit lines and 95% confidence intervals, respectively. Significant effects are indicated by *P <0.05, **P < 0.01, ***P < 0.001.

**Figure S4.**
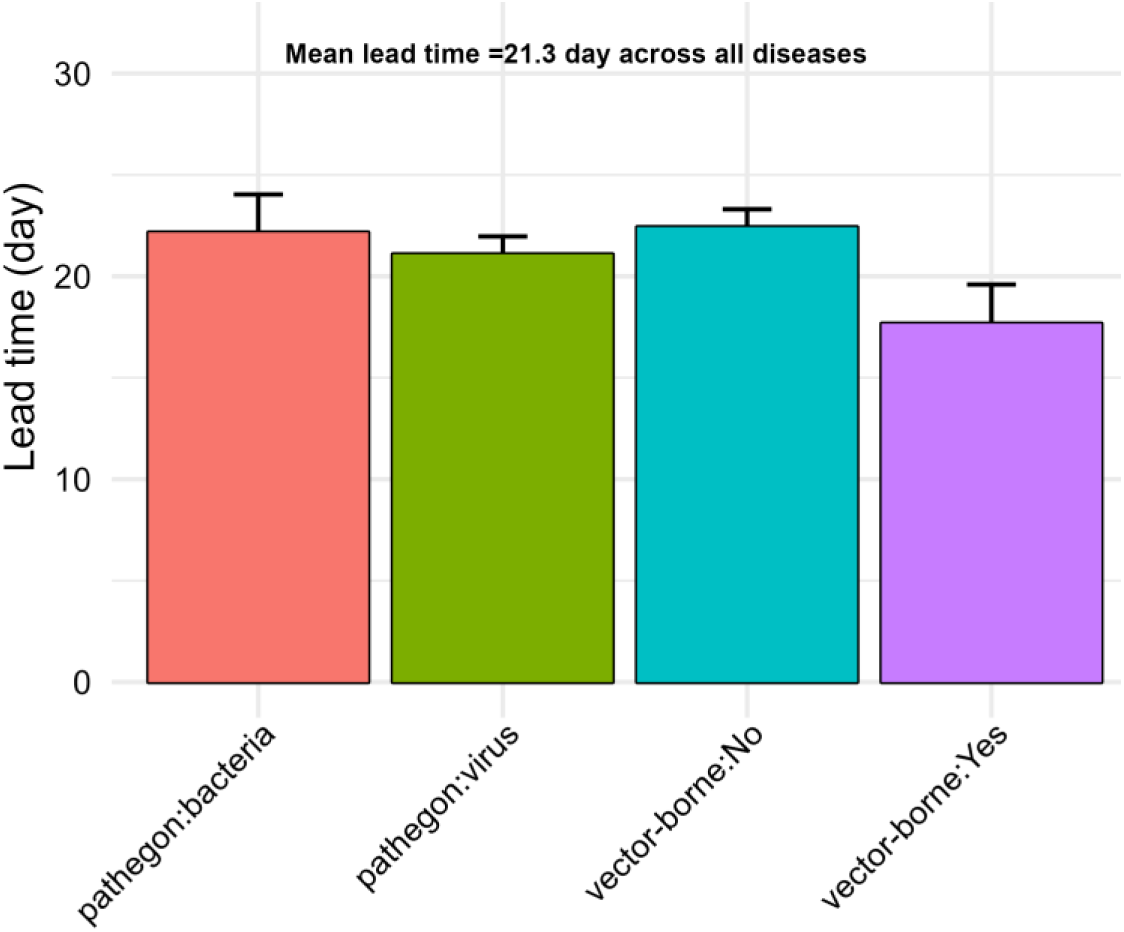
The lead time estimated from the time-to-event analysis based on daily data. The gray bar represents the mean, while the associated error bar illustrates the standard error. The lead time, i.e. days between the first instance that survival regression model was significant and the start of the outbreak, was computed by the best single RI (Standard Deviation, window length=10 days; Fig 3a-b, Fig S1). The predictor was the tau value (𝜏) of the RIs. The response variable was a time series of 0 values indicating non-outbreak days (𝑅_𝑒_ < 1) and a single 1 value indicating the hypothetical time point at which the outbreak could be anticipated in advance.

**Figure S5.**
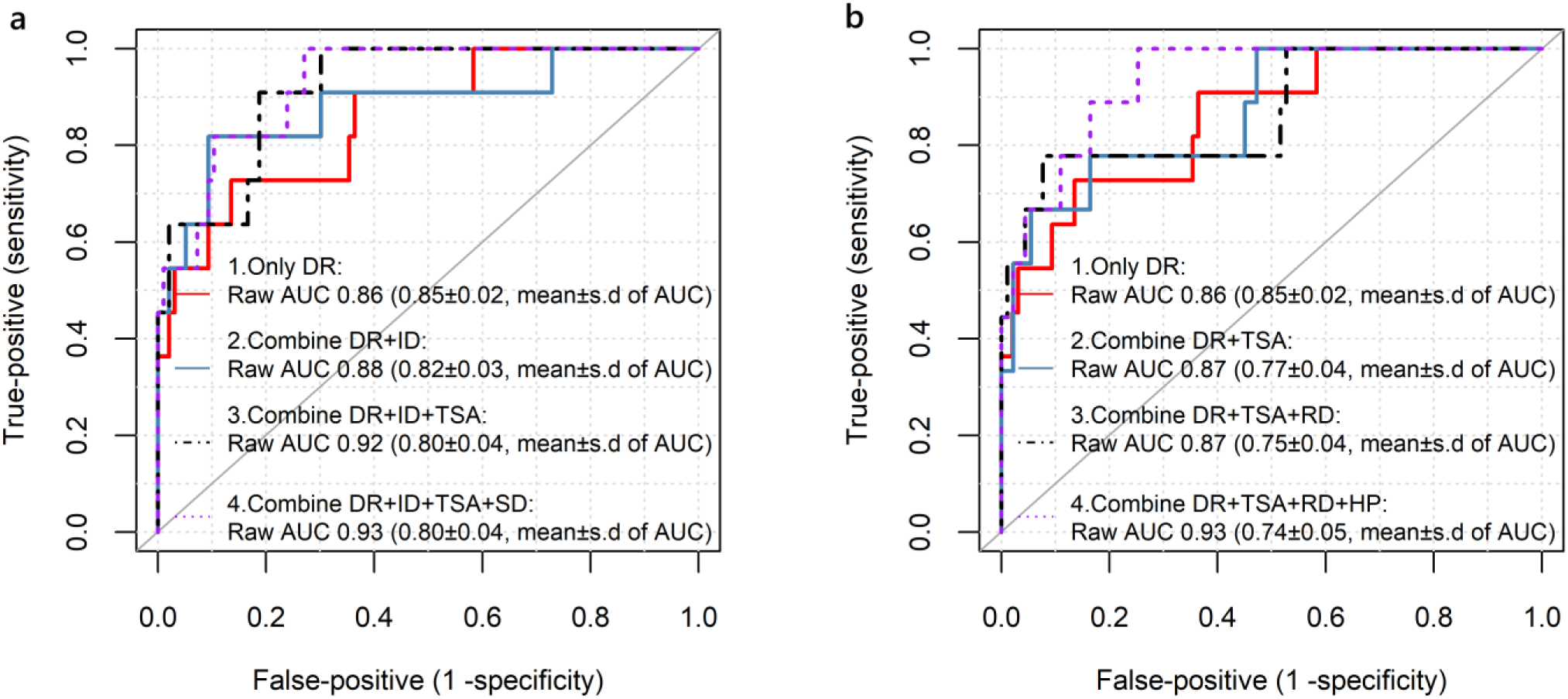
For the aggregated weekly data, the accuracy of resilience indicators (RIs) as early warning signals (EWSs) of outbreaks across 31 diseases, examined using mean area under the receiver operating characteristic curves (AUC) and cross validation. All RIs used the window length (i.e. 4 weeks) that maximized the AUC for the most accurate RI (i.e. Density ratio; mean AUC=0.85) (**a-b**) across all RIs and window lengths. We sequentially added either the next best performing single RI (**a**) or the next most orthogonal RI (i.e., least correlated based on Principal Component Analysis) to the best performing RI (**b**) until we tested up to four total RIs in our models. Importantly, sequentially adding the next most orthogonal RI to the model ensures that the baseline AUC goes up with each RI addition, but it does not guarantee that the mean AUC from the cross validation will continually rise with each added RI. Density ratio: DR, index of dispersion: ID, time series acceleration: TSA, standard deviation: SD, Relative dispersion: RD, Hurst exponent: HP. The AUC values in parenthesis (**a-b**) represent the mean and s.d from cross-validation.

**Figure S6.**
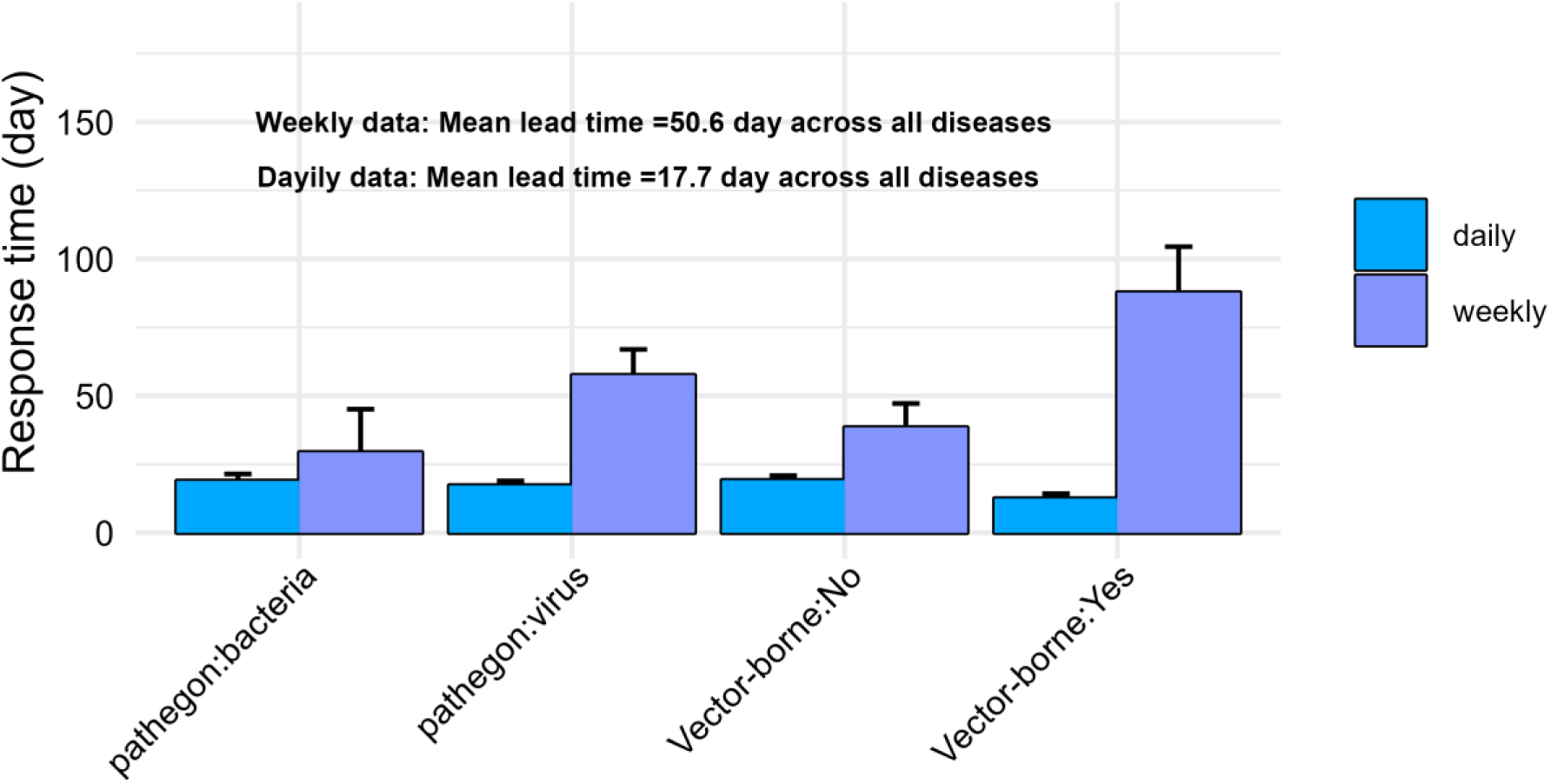
The lead time estimated from RIs approach based on daily and coarse- scale weekly data. The weekly data was derived by aggregating daily data to a weekly scale through weekly averaging. Apart from differences in data resolution, all other methods for calculating lead time for weekly data are identical to those applied to daily data using the RIs approach (see Methods). The gray bar represents the average, while the associated error bar illustrates the standard error. The lead time for weekly data was computed by the best performing single RI (Density ratio, window length=4 weeks; Fig. S5). For the daily data, the lead time was computed by the best performing single RI (Standard Deviation, window length=10 days; Fig3 & Fig S1).

**Figure S7.**
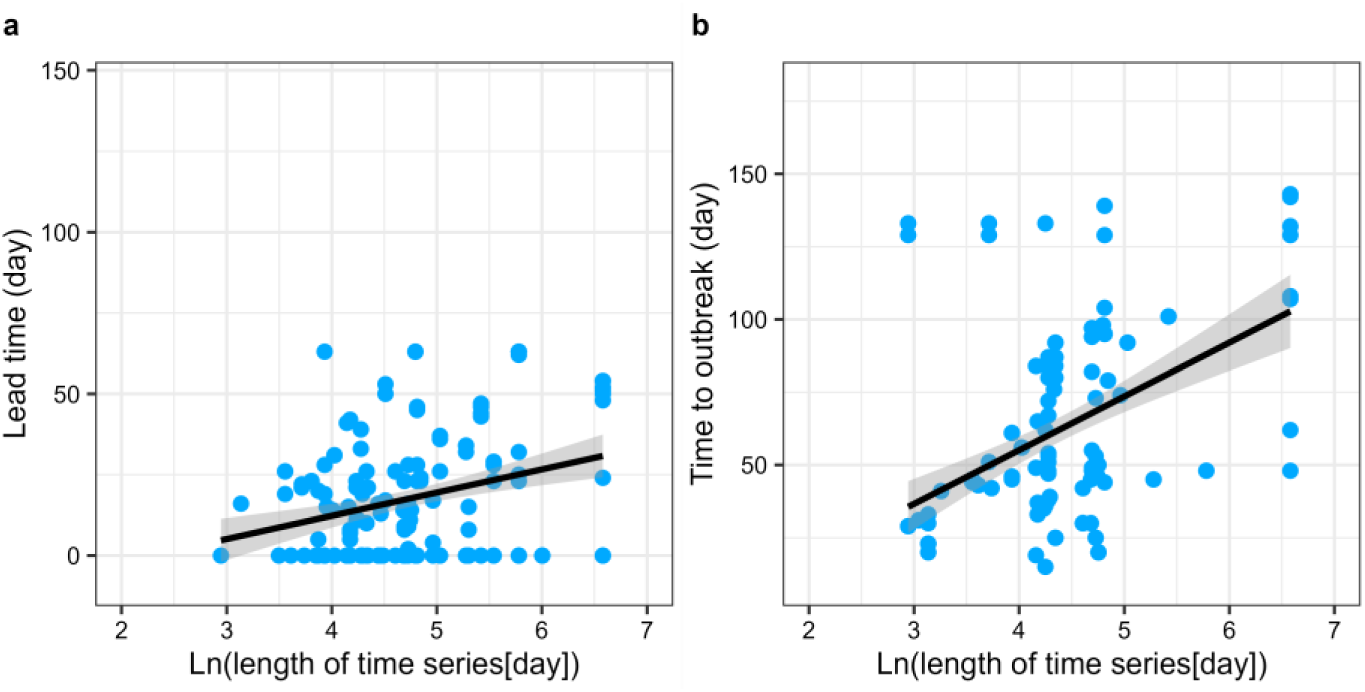
Effects of time series length on (a) lead time and (b) time to outbreak occurrence. Statistical results are presented in Table S1 and S3. The bold lines represent the significant best-fit trendlines, while the gray bands denote the corresponding 95% confidence bands. For the effects of the remaining variables on these response variables, refer to Figure 4 and 5.

**Figure S8.**
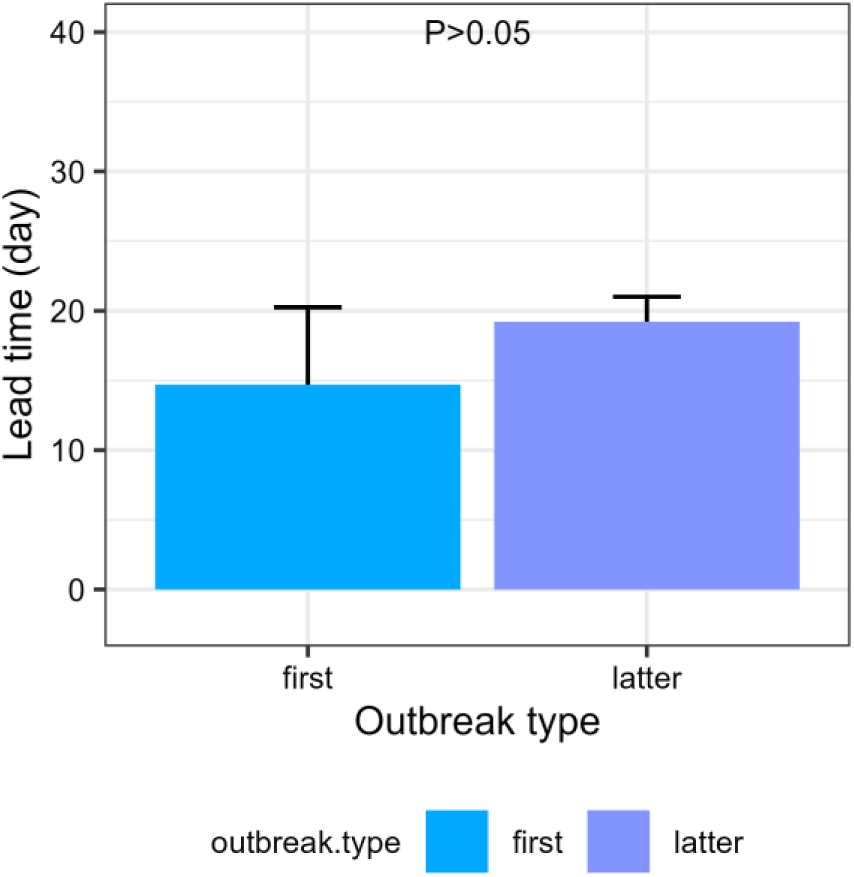
The lead time in responses to outbreak type. The lead time (mean ±s.e.) between the first outbreaks and subsequent outbreaks in a time series had no difference. Statistical results can be found in Table S1.

**Figure S9.**
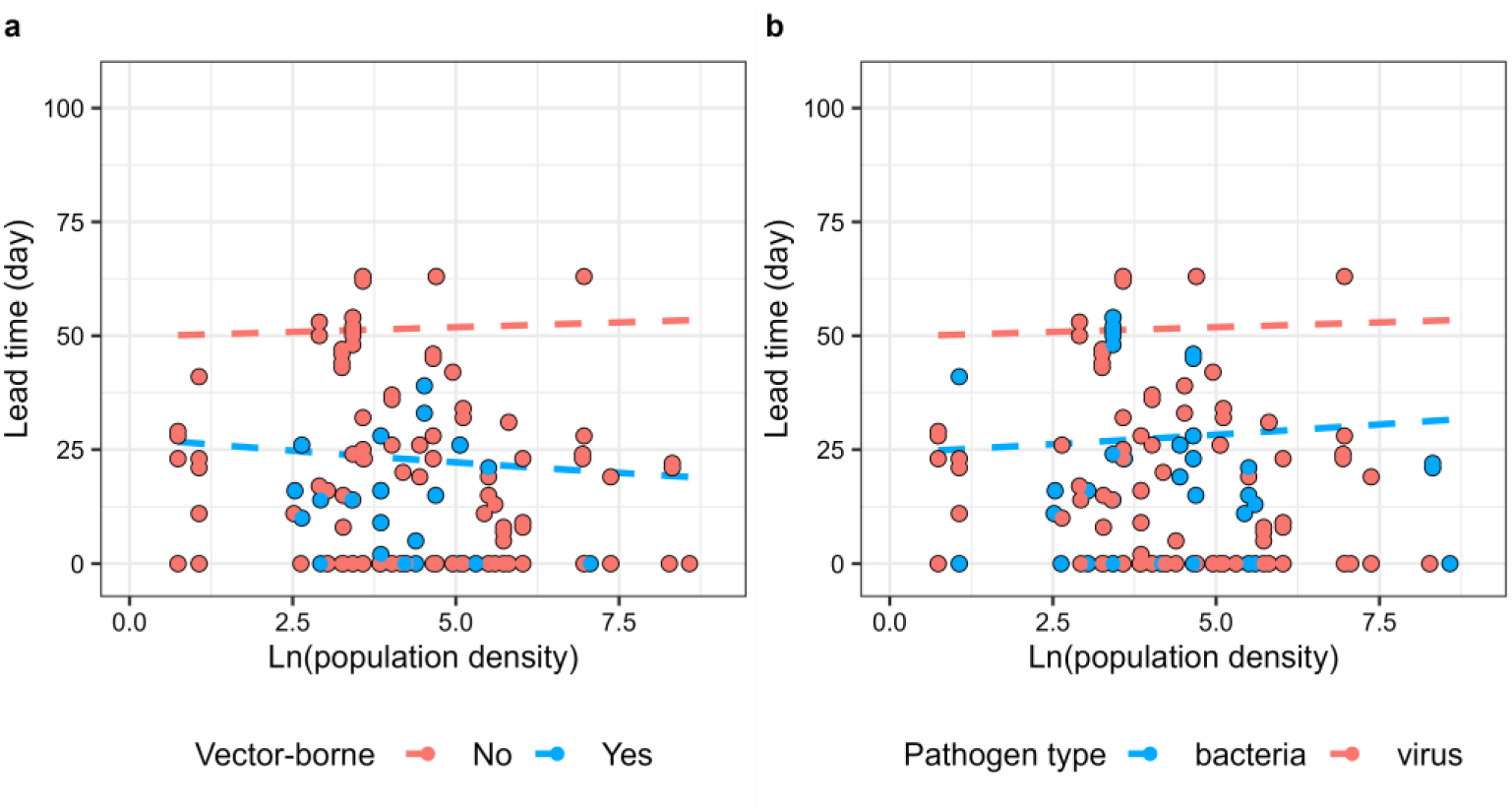
Effects of human population density (a, b) on the amount of lead time before disease outbreaks across 31 diseases. Dashed lines indicate non-significant trendlines. Details of statistical results are available in Table S1. For the effects of the remaining variables—temperature, precipitation, HDI, pathogen incubation time, and two parasite- pathogen traits—on lead time, refer to Figure 4.

**Figure S10.**
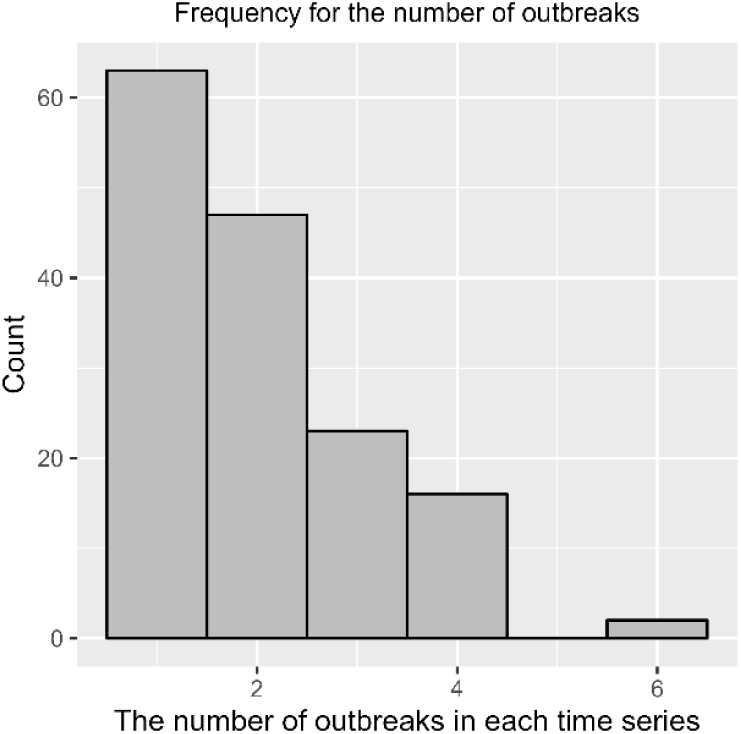
Frequency of the number of outbreaks per time series with the average number of secondary infections caused by a single infected individual at a given time point (𝑹_𝒆_ ≥ 1).

### Supplementary Information 1

The method developed by Cori et al. (*33*) defined the 𝑅_𝑒_ as the ratio of the number of new infections at time t (𝐼_𝑡_) to the total infectiousness of individuals up to that point.

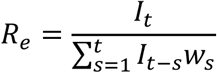

The total infectiousness is expressed as ∑^𝑡^ 𝐼_𝑡−𝑠_𝑤_𝑠_, representing the sum of infection incidences from previous time steps (up to t-1), weighted by 𝑤_𝑠_. The 𝑤_𝑠_ is a probability distribution (hence summing to 1) describing the average infectiousness profile after infection, which can be directly estimated using Markov Chain Monte Carlo (dic.fit.mcmc function in coarseDataTools package) (*48*). To account for the uncertainty in the distribution of 𝑤_𝑠_, we assume that 𝑤_𝑠_ follows a Gamma distribution, with its mean (𝜇) and standard deviation (𝜎) varying according to truncated normal distributions (*33*). Following the methodology of Cori et al. (*33*), we generate 1,000 pairs of mean and standard deviation, denoted as (𝜎, 𝜇)^1^, … ., (𝜎, 𝜇)^1000^, with the constraint that 𝜎 < 𝜇 in each pair. This constraint ensures that the Gamma probability density function of 𝑤_𝑠_ is null at t=0. For each pair (𝜎, 𝜇)^𝑘^, we draw 1,000 realizations of 𝑅_𝑒_ from its posterior distribution, conditional on the corresponding (𝜎, 𝜇)^𝑘^. This process yields a total sample size of 1,000,000 (1,000 × 1,000) from the joint posterior distribution of 𝑅_𝑒_. The posterior mean, standard deviation, and the 0.025, 0.05, 0.25, 0.5, 0.75, 0.95, and 0.975 quantiles of 𝑅_𝑒_ are eventually obtained from this sample.

Unlike methods that require fitting mechanistic dynamic models to incidence or case data for estimating the 𝑅_𝑒_, the approaches developed by Cori et al. (*33*), along with the other two approaches developed by Wallinga and Teunis (*49*), and Bettencourt and Ribeiro (*50*), are generic and commonly used model-free method that rely on case incidence data for 𝑅_𝑒_ estimation. However, among the three, the method by Cori et al. (*33*) had higher accuracy and addressed the limitations present in the other two approaches. First, Cori et al. (*33*) uses data from before time *t*, while Wallinga and Teunis (*49*) require data from after time *t* (*33*). This reliance makes the latter approach conceptually and methodologically less suitable for near real-time estimation, particularly when the objective is to assess the impact of interventions or other external factors on transmission dynamics at a specific point in time (*33*, *34*). Second, the method proposed by Cori et al. (*33*) can accommodate any positive, discrete generation interval distribution. In contrast, the approach by Bettencourt and Ribeiro (*50*) implicitly assumes that the generation interval follows an exponential distribution-as it is derived from a Susceptible-Infectious-Recovered (SIR) model, which restricts the applicability of the latter method (*33*, *34*). Third, the method developed by Cori et al. (*33*) demonstrates greater robustness when applied to data with small aggregation time steps (e.g., daily data), enabling the generation of smoother estimates (*34*).

### Supplementary Information 2

#### Details of the 17 resilience indicators

1. *Standard deviation, Coefficient of Variation, Index of Dispersion and First differenced variance*.

As a system approaches a critical transition, the slowing rate of return to equilibrium can result in greater fluctuations facing disturbances (*36*). Large disturbances can push the system beyond the boundaries of alternative stable states—a phenomenon known as flickering. Both critical slowing down and flickering contribute to an increase in variance prior to a full transition, making it a key indicator of early warning (*36*).

Variance can be measured using standard deviation SD 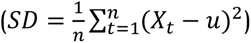 (36), the coefficient of variation 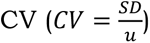 (36), index of dispersion ID (*36*), or first differenced variance (FDV) (*12*), The rise in variance reflects the system’s diminishing ability to recover from perturbations, serving as a useful diagnostic for detecting impending state shifts.

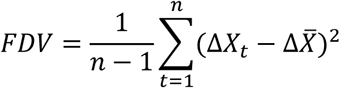

Where Δ𝑋_𝑡_ is the difference among disease cases at time 𝑡 and 𝑡 − 1, e.g. Δ𝑋_𝑡_ = 𝑋_𝑡_ − 𝑋_𝑡−1_. Δ𝑋̅ is the mean of these difference.

1. *Skewness, Kurtosis and Density Ratio*.

Certain types of disturbances can drive the system state toward the boundary between two alternative stable states (*36*). At these boundaries, the system dynamics slow down, leading to an increased skewness (SK) of a time series, i.e. an asymmetric in distribution of observed values (*36*). Skewness is defined as the standardized third moment around the mean, capturing the degree of asymmetry in a distribution 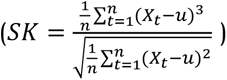 (36).

In addition, disturbances and flickering can push the system state toward more extreme values as it approaches a critical transition (*36*). These effects lead to an increased frequency of rare events, resulting in fatter tails in the time series distribution. Consequently, the kurtosis (KU) of the time series can rise prior to the transition, reflecting an elevated occurrence of extreme deviations from the mean. Kurtosis is defined as the standardized fourth moment around the mean, capturing the heaviness of the tails in a distribution 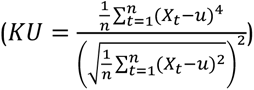 (36).

As a system approaches a critical transition, this also results in changes in the probability density function of the system’s states over time (*36*). The Density Ratio (DR) can serve as an early warning signal by identifying shifts in the probability distribution. The DR measures these changes by comparing the probability density of recent observations (𝑃_𝑝𝑟𝑒𝑠𝑒𝑛𝑡_) to a reference (baseline) distribution from a stable period (𝑃_𝑟𝑒𝑓_) (*36*), expressed as 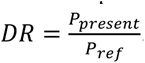.

1. *Autocorrelation at lag 1, 2 and 3, and Autocovariance*. As a system nears a critical transition, the rate of returns to equilibrium following a small perturbation decreases (*36*). This slow rate of return is termed as critical slowing down (*36*). As the rate of return decreases, the state of the system becomes increasingly similar between consecutive observations, where the time series exhibits an increase of autocorrelation (*36*). Therefore, autocorrelation at low lags can serve as warning signals of an impending transition (*36*).

Autocorrelation at lag 1 can be measured by a conditional least-squares method to fit an autoregressive model of order 1 (AR(1)), which is expressed as:

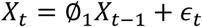

𝑋_𝑡_ and 𝑋_𝑡−1_ are the disease cases at time t and t-1, respectively. 𝜖_𝑡_ is a Gaussian white noise error term with mean zero and constant variance. ∅_1_ is the autoregressive coefficient and serves as a proxy for Autocorrelation at lag 1 (*36*). Similarly, the Autocorrelation at lag 2 and 3 can be expressed as 𝑋_𝑡_ = ∅_2_𝑋_𝑡−2_ + 𝜖_𝑡_ and 𝑋_𝑡_ = ∅_3_𝑋_𝑡−3_ + 𝜖_𝑡_, respectively. Here ∅_2_ and ∅_3_ are Autocorrelation at 2 and 3, respectively.

In addition, as a system nears a transition, the decline in the rate of return causes deviations from equilibrium for longer periods. This prolonged persistence increases the dependence between observed values in the time series, resulting in elevated autocovariance (AU)(*10*, *51*). Autocovariance can thus serve as a metric for quantifying the persistence of fluctuations 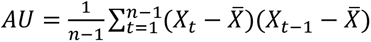 (10, 51)

1. *Wavelet filtering and Wavelet reddening*.

When disease transmission is subject to seasonal forcing, wavelet-based warning signals can capture the temporal variability of disease dynamics and serve as robust indicators of impending epidemiological changes (*52*).

As Miller et al (*52*), Wavelet filtered reddening 𝑊𝐹 is defined as any quantity proportional to the variance of the time series after it is filtered to include only its components in a wavelet transform’s frequency domain, calculated as 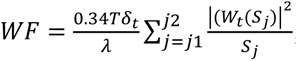, where 𝑇 is length of the time series, 𝛿_𝑡_ is the difference in time between observations, 𝜆 represents the ratio of the Fourier period of the wavelet function to its corresponding scale, 𝑆_𝑗_ is a wavelet scale and 𝑊_𝑡_(𝑆_𝑗_) represents the wavelet transform of the observed disease time series, localized at time index 𝑡 and the wavelet scale 𝑆_𝑗_ (*52*). All parameters within the equation for 𝑊𝐹 was estimated in accordance with the methodology of Miller et al. (*52*).

Next, wavelet spectral reddening (WR) is defined as the median scale of the wavelet spectral density at a given time 𝑡, expressed as 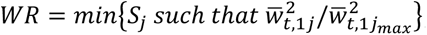, where the wavelet scale increases from 1 to 𝑗_𝑚𝑎𝑥_ (*52*).

1. *Hurst exponent, Relative dispersion, and Time series acceleration*.

As a system approaches critical slowing down, the *Hurst exponent* (HE) is expected to increase, and it has been demonstrated to effectively predict the transition in advance of the tipping point (*53*, *54*). HE is calculated by the algorithm of Multifractal detrended fluctuation analysis, where the HE is computed as the linear slope for the variation of fluctuations 𝐹^2^against the span size 𝑤 (*53*). Here, the variation of fluctuations 𝐹^2^ is determined as 𝐹^2^ = 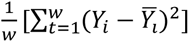, where 𝑌 the cumulative deviation series from the overall mean of the time series and expressed as 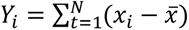 (*53*). The 𝑌̅_𝑖_ is the polynomial fit of order 1 against each of these segments 𝑤. The 𝑤 is to divide the entire time series into non-overlapping segments, and the 𝑤 varies from 2 to 𝑁 − 2, where the 𝑁 is the length of entire time series.

Following the Cerqueira et al. (*55*) and Wang et al. (*56*), *Relative Dispersion* is estimated as the ratio between the standard deviation of the time series and the standard deviation of the differenced time series. *Time series acceleration* the second derivative of the observed values with respect to time, which is expected to decrease as the system nears to tipping point (*55*).

1. *The absolute value of dominant eigenvalue (Max(eigen))*.

Theoretically, as a discreate time system nears a tipping point, the absolute value of dominant eigenvalue (Max(eigen)) from Jacobian Matrix approaches 1 (*57*). The elements in Jacobian matrix (𝑱) can be estimated by S-map, 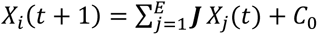, where 𝐸 is the embedding dimension consisting of number of 𝐸 state variables (*57*, *58*). The S-map algorithm constructs a local linear model (**C**) to predict the future state at time 𝑡 + 1 using the state vector 𝑋_𝑗_ from the reconstructed state space (*57*, *58*). However, unlike traditional linear regression models, the linear approximation in the S-map algorithm is performed locally, assigning greater weights to data points that are in close to the target state vector 𝑋_𝑗_ (*57*, *58*).

For the disease cases time series, the S-map algorithm reconstructs the Jacobian matrix (𝑱) using a single observed variable (i.e., disease cases). This follows the principle of lagged coordinate embedding (i.e., univariate embedding), which reconstructs the underlying system’s state space from time-delayed versions of the univariate time series (*57*). Since the reconstructed state space is a delay embedding of a univariate time series, the resulting Jacobian matrix (𝑱) is characterized by a sparse structure. Specifically, all elements of 𝑱 are zero except for the lower off-diagonal elements, which are equal to unity, and the first row, which contains the nontrivial linear approximation that governs the system’s local dynamics (*57*).

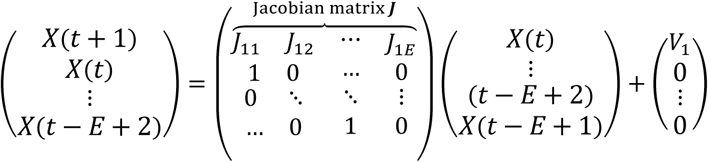

Since the Jacobian matrix (𝑱) is evaluated at each time point 𝑡 (i.e., the local Jacobian 𝑱_𝑡_), it exhibits time-varying dynamics. From these local Jacobians, the absolute value of the dominant eigenvalue *(Max(eigen))* can be computed at each time step, providing insights into the temporal changes of the disease system’s stability shifting from disease free to outbreak states (*57*). The best embedding dimension 𝐸 is selected by using the simplex projection that gave the lowest mean absolute error (*59*).

### Supplementary tables

**Table S1.**
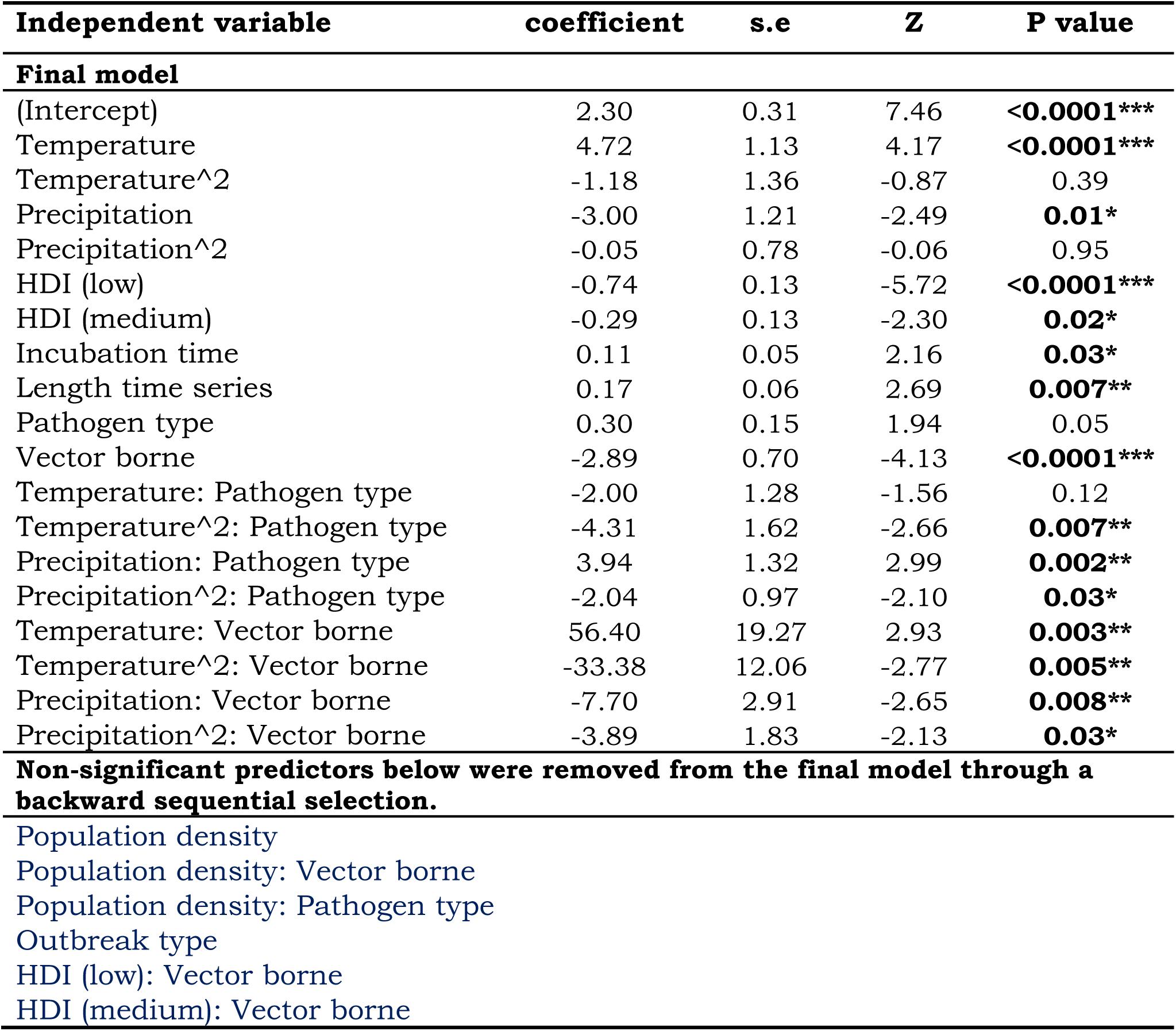
Results from a generalized linear mixed model examining the main effects of temperature, precipitation, human development index (HDI), pathogen incubation time, human population density, time series length, two pathogen traits (vector-borne or not; virus vs. bacterial pathogen), outbreak type (first or subsequent outbreak in a time series), and the pairwise interactions between the two pathogen traits and temperature, precipitation, HDI and human population density on lead time. In the model, we treated regions nested within disease types as random intercepts. The parasite taxa were limited to viruses and bacteria because our dataset had no fungi or helminths (Table S4) and the sample size for protozoa was too small to justify analysis (only two parasite species). A significant quadratic coefficient indicates a second-order polynomial function between a predictor and lead time. The blue text denotes non-significant predictors that were removed from the final model through a backward sequential selection. \*\*\**P*< 0.001, \*\**P*<0.01, \**P*<0.05. The model’s marginal R^2^ is 0.12, and the conditional R^2^ is 0.17. The model AIC value before and after removing the blue non-significant predictors was 1131.57 and 1121.83, respectively.

| Independent variable | coefficient | s.e | Z | P value |
| --- | --- | --- | --- | --- |
| **Final model** |  |  |  |  |
| (Intercept) | 2.30 | 0.31 | 7.46 | <0.0001*** |
| Temperature | 4.72 | 1.13 | 4.17 | <0.0001*** |
| Temperature^2 | -1.18 | 1.36 | -0.87 | 0.39 |
| Precipitation | -3.00 | 1.21 | -2.49 | 0.01* |
| Precipitation^2 | -0.05 | 0.78 | -0.06 | 0.95 |
| HDI (low) | -0.74 | 0.13 | -5.72 | <0.0001*** |
| HDI (medium) | -0.29 | 0.13 | -2.30 | 0.02* |
| Incubation time | 0.11 | 0.05 | 2.16 | 0.03* |
| Length time series | 0.17 | 0.06 | 2.69 | 0.007** |
| Pathogen type | 0.30 | 0.15 | 1.94 | 0.05 |
| Vector borne | -2.89 | 0.70 | -4.13 | <0.0001*** |
| Temperature: Pathogen type | -2.00 | 1.28 | -1.56 | 0.12 |
| Temperature^2: Pathogen type | -4.31 | 1.62 | -2.66 | 0.007** |
| Precipitation: Pathogen type | 3.94 | 1.32 | 2.99 | 0.002** |
| Precipitation^2: Pathogen type | -2.04 | 0.97 | -2.10 | 0.03* |
| Temperature: Vector borne | 56.40 | 19.27 | 2.93 | 0.003** |
| Temperature^2: Vector borne | -33.38 | 12.06 | -2.77 | 0.005** |
| Precipitation: Vector borne | -7.70 | 2.91 | -2.65 | 0.008** |
| Precipitation^2: Vector borne | -3.89 | 1.83 | -2.13 | 0.03* |
| **Non-significant predictors below were removed from the final model through a backward sequential selection.** |  |  |  |  |
| Population density |  |  |  |  |
| Population density: Vector borne |  |  |  |  |
| Population density: Pathogen type |  |  |  |  |
| Outbreak type |  |  |  |  |
| HDI (low): Vector borne |  |  |  |  |
| HDI (medium): Vector borne |  |  |  |  |

**Table S2.** Results from a mixed effect Cox proportional hazard model examining the main effects of temperature, precipitation, human development index (HDI), pathogen incubation time, human population density, time series length, two pathogen traits (vector-borne or not; virus vs. bacterial pathogen), outbreak type (first or subsequent outbreak in a time series) and the pairwise interactions between the two pathogen traits and temperature, precipitation, HDI and human population density on the timing of outbreak occurrence. In the model, we treated regions nested within disease types as random intercepts. The parasite taxa were limited to viruses and bacteria because our dataset had no fungi or helminths (Table S4) and the sample size for protozoa was too small to justify analysis (only two parasite species). A positive coefficient signifies an increased hazard of outbreak occurrence, and consequently a reduced time until an outbreak. A significant quadratic coefficient indicates a second-order polynomial function between a predictor and hazard of outbreak occurrence. The blue text denotes non-significant predictors that were removed from the final model through a backward sequential selection. Model’s Nagelkerke R² is 0.29. \*\*\**P*< 0.001, \*\**P*<0.01, \**P*<0.05. The model AIC value before and after removing the blue non-significant predictors was 536.73 and 520.35, respectively.

| Independent variable | coefficient | s.e | Z | P value |
| --- | --- | --- | --- | --- |
| **Final model** |  |  |  |  |
| Temperature^2 | 7.36 | 1.44 | 5.10 | **<0.0001\*\*\*** |
| Temperature | -1.29 | 1.44 | -0.90 | 0.36 |
| Precipitation^2 | 2.97 | 0.67 | 4.43 | **<0.0001\*\*\*** |
| Precipitation | 0.14 | 0.76 | 0.18 | 0.85 |
| HDI (low) | 0.95 | 2.09 | 0.46 | 0.64 |
| HDI (medium) | 0.12 | 2.04 | 0.06 | 0.95 |
| Length time series | -4.21 | 0.83 | -5.05 | **<0.0001\*\*\*** |
| Pathogen type | 5.06 | 1.94 | 2.60 | **0.009\*\*** |
| Vector borne | -0.45 | 2.42 | -0.19 | 0.85 |
| Temperature^2: Pathogen type | -8.01 | 1.56 | -5.15 | **<0.0001\*\*\*** |
| Temperature: Pathogen type | -3.20 | 1.70 | -1.88 | 0.06 |
| Precipitation: Vector borne | -5.43 | 1.05 | -5.18 | **<0.0001\*\*\*** |
| Precipitation^2: Vector borne | -2.30 | 0.93 | -2.47 | **0.01\*** |
| HDI (low): Vector borne | 14.60 | 4.09 | 3.57 | **0.0003\*\*\*** |
| HDI (medium): Vector borne | 12.29 | 3.92 | 3.14 | **0.001\*\*** |
| **Non-significant predictors below were removed from the final model through a backward sequential selection.** |  |  |  |  |
| Population density |  |  |  |  |
| Population density: Pathogen type |  |  |  |  |
| Population density: Vector borne |  |  |  |  |
| Incubation time |  |  |  |  |
| Outbreak type |  |  |  |  |
| Precipitation: Pathogen type |  |  |  |  |
| Precipitation^2: Pathogen type |  |  |  |  |
| Temperature^2: Vector borne |  |  |  |  |
| Temperature: Vector borne |  |  |  |  |

**Table S3.** Given that there is currently no reliable way to visualize the results of a Cox proportional hazards (coxme) model, we replicated the final coxme model (Table S2) as a generalized linear mixed model (glmmTMB function in glmmTMB package) with time to an outbreak as a negative binomial dependent variable, shown below. Given that the statistical outcomes of the two models had verified direction (i.e. a positive coefficient in Table S2 signifies an increased hazard of outbreak occurrence and consequently a reduced time until an outbreak in Table S3, and vice versa) and similar significance levels, we used the glmmTMB to generate figures using the ggpredict function in the ggeffects R package.

| Independent variable | coefficient | s.e | Z | P value |
| --- | --- | --- | --- | --- |
| Temperature^2 | -1.27 | 0.46 | 2.75 | **0.006\*\*** |
| Temperature | 3.73 | 0.62 | 6.03 | 0.06 |
| Precipitation^2 | -3.63 | 0.62 | -5.85 | **<0.0001\*\*\*** |
| Precipitation | -0.26 | 0.63 | -0.42 | 0.67 |
| HDI (low) | -0.03 | 0.11 | -0.27 | 0.79 |
| HDI (medium) | -0.13 | 0.12 | 1.15 | 0.25 |
| Length time series | 0.21 | 0.04 | 4.77 | **<0.0001\*\*\*** |
| Pathogen type | -1.10 | 0.08 | -1.14 | **0.04\*** |
| Vector borne | 0.11 | 0.18 | -0.61 | 0.54 |
| Temperature^2: Pathogen type | 3.07 | 1.12 | -2.68 | **0.007\*\*** |
| Temperature: Pathogen type | 2.32 | 1.05 | -2.12 | 0.05 |
| Precipitation: Vector borne | 3.12 | 1.26 | 2.46 | **0.01\*** |
| Precipitation^2: Vector borne | 1.84 | 0.91 | 2.05 | **0.04\*** |
| HDI (low): Vector borne | -0.73 | 0.22 | -3.40 | **0.0006\*\*\*** |
| HDI (medium): Vector borne | -0.14 | 0.25 | 0.15 | **0.03\*** |

**Table S4.**
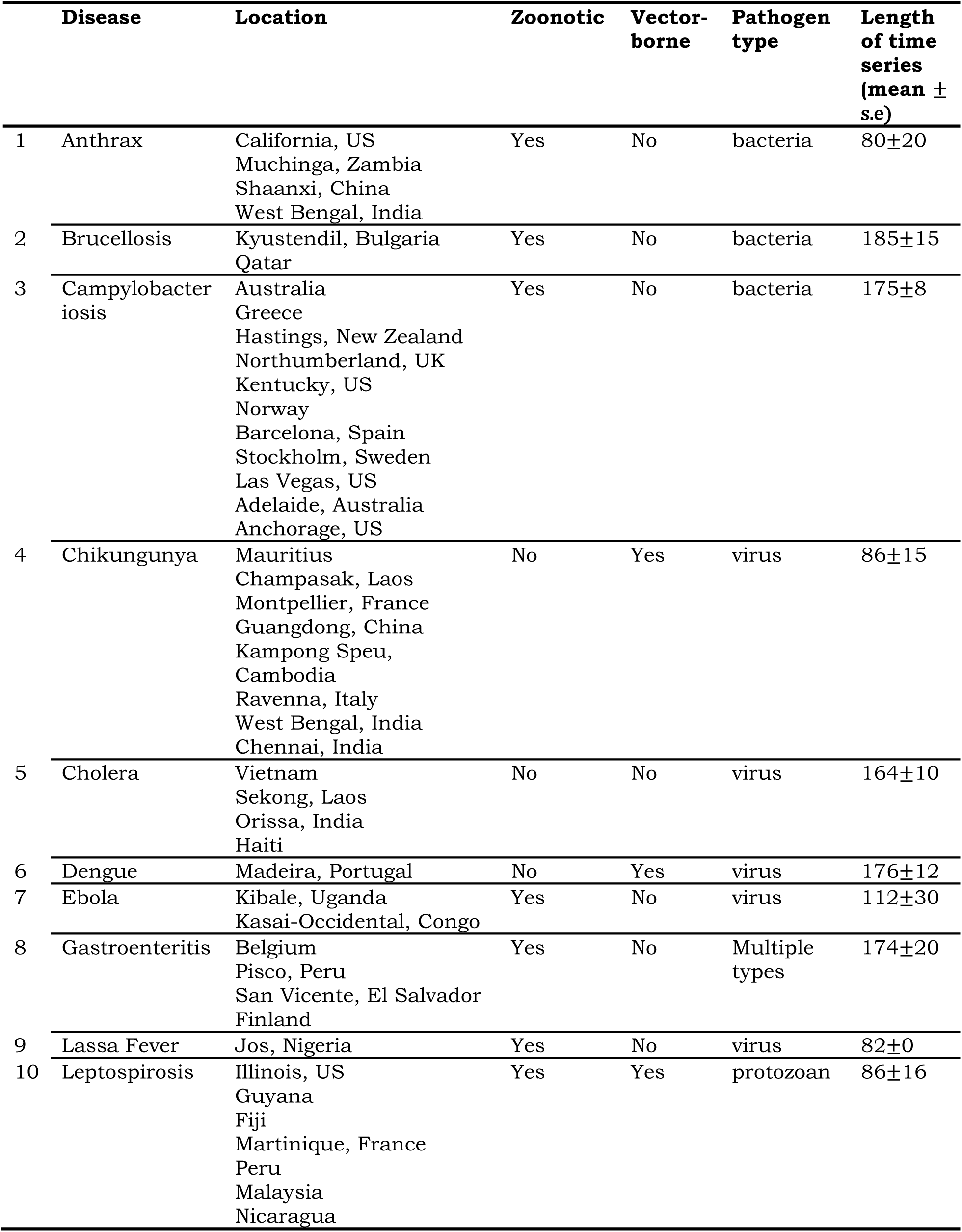
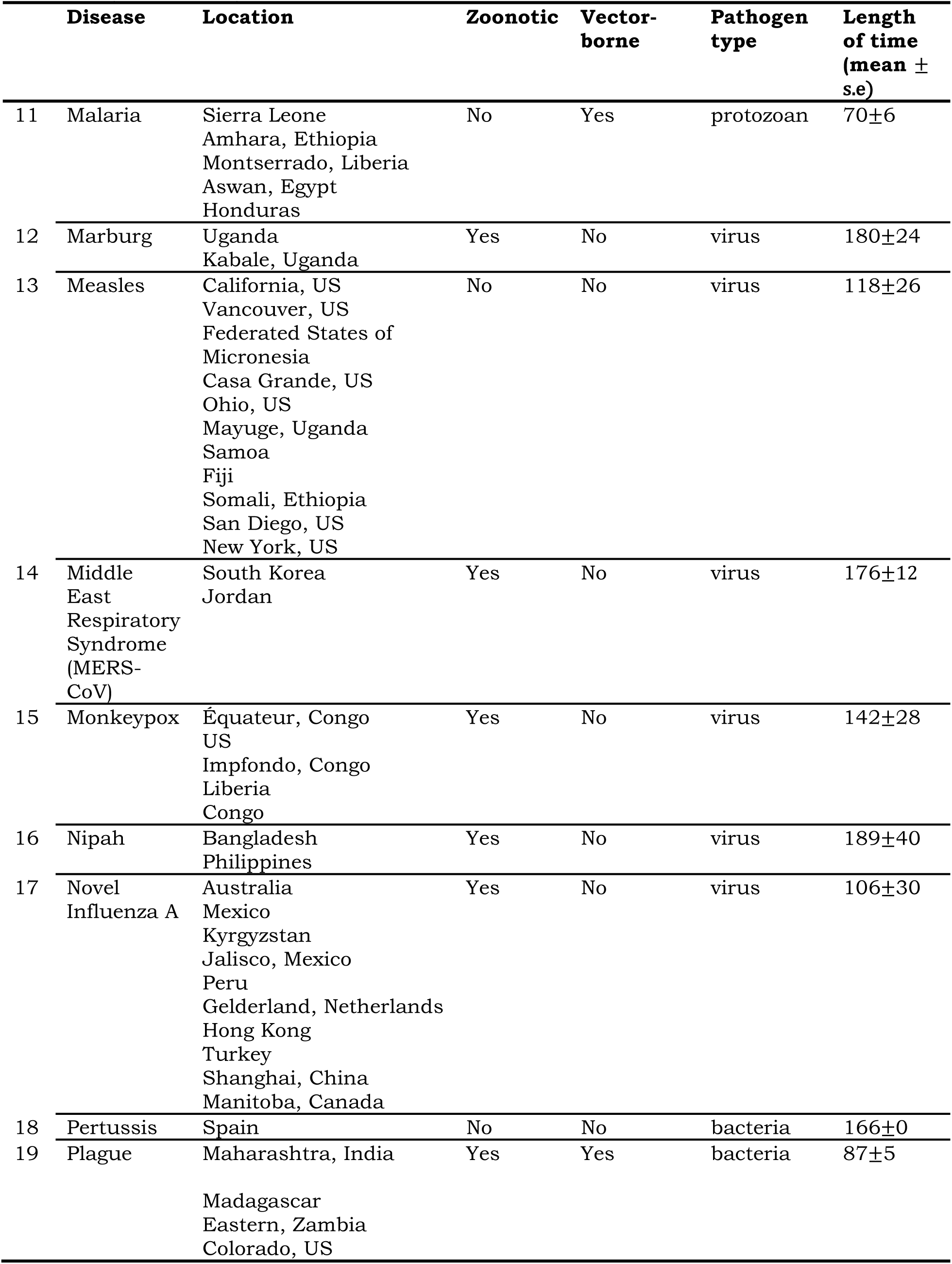
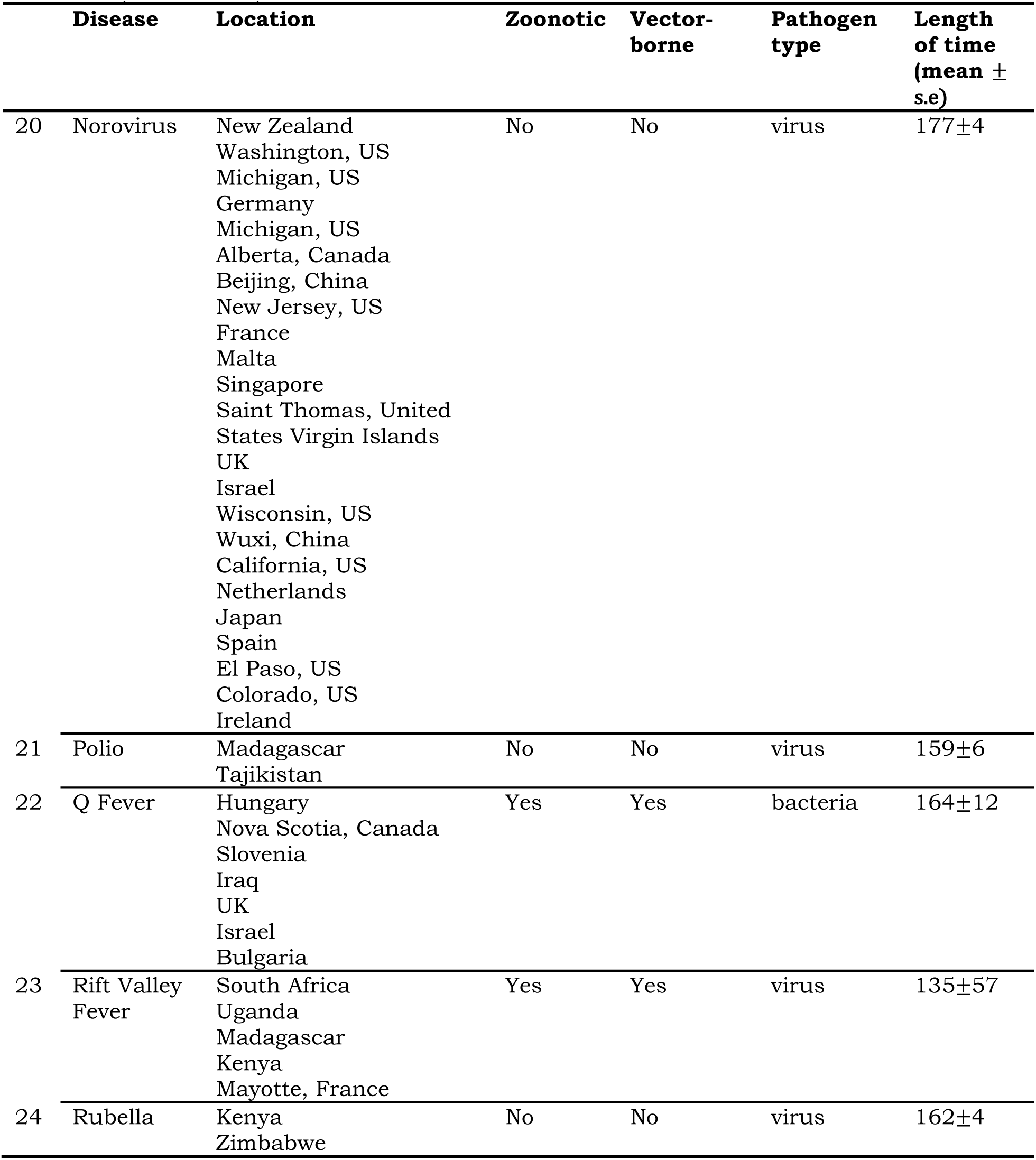
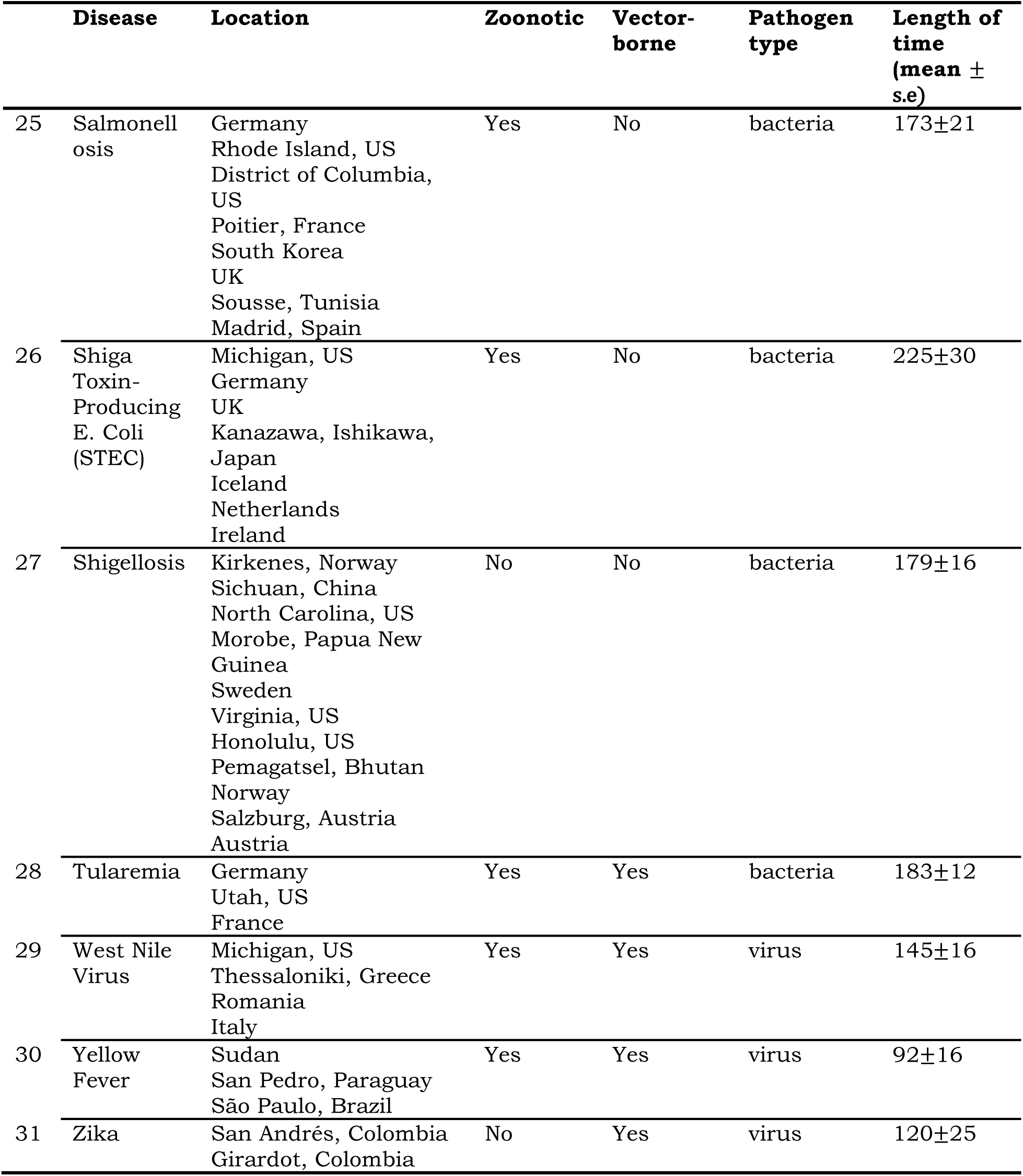
All 31 diseases used in the analyses, their corresponding regions, and traits of their pathogens and time series.

|  | **Disease** | **Location** | **Zoonotic** | **Vector-**<br>**borne** | **Pathogen**<br>**type** | **Length**<br>**of time**<br>**series**<br>**(mean ±**<br>**s.e)** |
| --- | --- | --- | --- | --- | --- | --- |
| 1 | Anthrax | California, US<br>Muchinga, Zambia<br>Shaanxi, China<br>West Bengal, India | Yes | No | bacteria | 80±20 |
| 2 | Brucellosis | Kyustendil, Bulgaria<br>Qatar | Yes | No | bacteria | 185±15 |
| 3 | Campylobacter<br>iosis | Australia<br>Greece<br>Hastings, New Zealand<br>Northumberland, UK<br>Kentucky, US<br>Norway<br>Barcelona, Spain<br>Stockholm, Sweden<br>Las Vegas, US<br>Adelaide, Australia<br>Anchorage, US | Yes | No | bacteria | 175±8 |
| 4 | Chikungunya | Mauritius<br>Champasak, Laos<br>Montpellier, France<br>Guangdong, China<br>Kampong Speu,<br>Cambodia<br>Ravenna, Italy<br>West Bengal, India<br>Chennai, India | No | Yes | virus | 86±15 |
| 5 | Cholera | Vietnam<br>Sekong, Laos<br>Orissa, India<br>Haiti | No | No | virus | 164±10 |
| 6 | Dengue | Madeira, Portugal | No | Yes | virus | 176±12 |
| 7 | Ebola | Kibale, Uganda<br>Kasai-Occidental, Congo | Yes | No | virus | 112±30 |
| 8 | Gastroenteritis | Belgium<br>Pisco, Peru<br>San Vicente, El Salvador<br>Finland | Yes | No | Multiple<br>types | 174±20 |
| 9 | Lassa Fever | Jos, Nigeria | Yes | No | virus | 82±0 |
| 10 | Leptospirosis | Illinois, US<br>Guyana<br>Fiji<br>Martinique, France<br>Peru<br>Malaysia<br>Nicaragua | Yes | Yes | protozoan | 86±16 |

**Table S4 (continued).**
|  | **Disease** | **Location** | **Zoonotic** | **Vector-**<br>**borne** | **Pathogen**<br>**type** | **Length**<br>**of time**<br>**(mean ±**<br>**s.e)** |
| --- | --- | --- | --- | --- | --- | --- |
| 11 | Malaria | Sierra Leone<br>Amhara, Ethiopia<br>Montserrado, Liberia<br>Aswan, Egypt<br>Honduras | No | Yes | protozoan | 70±6 |
| 12 | Marburg | Uganda<br>Kabale, Uganda | Yes | No | virus | 180±24 |
| 13 | Measles | California, US<br>Vancouver, US<br>Federated States of<br>Micronesia<br>Casa Grande, US<br>Ohio, US<br>Mayuge, Uganda<br>Samoa<br>Fiji<br>Somali, Ethiopia<br>San Diego, US<br>New York, US | No | No | virus | 118±26 |
| 14 | Middle<br>East<br>Respiratory<br>Syndrome<br>(MERS-CoV) | South Korea<br>Jordan | Yes | No | virus | 176±12 |
| 15 | Monkeypox | Équateur, Congo<br>US<br>Impfondo, Congo<br>Liberia<br>Congo | Yes | No | virus | 142±28 |
| 16 | Nipah | Bangladesh<br>Philippines | Yes | No | virus | 189±40 |
| 17 | Novel<br>Influenza A | Australia<br>Mexico<br>Kyrgyzstan<br>Jalisco, Mexico<br>Peru<br>Gelderland, Netherlands<br>Hong Kong<br>Turkey<br>Shanghai, China<br>Manitoba, Canada | Yes | No | virus | 106±30 |
| 18 | Pertussis | Spain | No | No | bacteria | 166±0 |
| 19 | Plague | Maharashtra, India<br>Madagascar<br>Eastern, Zambia<br>Colorado, US | Yes | Yes | bacteria | 87±5 |

**Table S4 (continued).**
|  | **Disease** | **Location** | **Zoonotic** | **Vector-**<br>**borne** | **Pathogen**<br>**type** | **Length**<br>**of time**<br>**(mean ±**<br>**s.e)** |
| --- | --- | --- | --- | --- | --- | --- |
| 20 | Norovirus | New Zealand<br>Washington, US<br>Michigan, US<br>Germany<br>Michigan, US<br>Alberta, Canada<br>Beijing, China<br>New Jersey, US<br>France<br>Malta<br>Singapore<br>Saint Thomas, United<br>States Virgin Islands<br>UK<br>Israel<br>Wisconsin, US<br>Wuxi, China<br>California, US<br>Netherlands<br>Japan<br>Spain<br>El Paso, US<br>Colorado, US<br>Ireland | No | No | virus | 177±4 |
| 21 | Polio | Madagascar<br>Tajikistan | No | No | virus | 159±6 |
| 22 | Q Fever | Hungary<br>Nova Scotia, Canada<br>Slovenia<br>Iraq<br>UK<br>Israel<br>Bulgaria | Yes | Yes | bacteria | 164±12 |
| 23 | Rift Valley<br>Fever | South Africa<br>Uganda<br>Madagascar<br>Kenya<br>Mayotte, France | Yes | Yes | virus | 135±57 |
| 24 | Rubella | Kenya<br>Zimbabwe | No | No | virus | 162±4 |

**Table S4 (continued).**
|  | **Disease** | **Location** | **Zoonotic** | **Vector-**<br>**borne** | **Pathogen**<br>**type** | **Length of**<br>**time**<br>**(mean ±**<br>**s.e)** |
| --- | --- | --- | --- | --- | --- | --- |
| 25 | Salmonellosis | Germany<br>Rhode Island, US<br>District of Columbia, US<br>Poitier, France<br>South Korea<br>UK<br>Sousse, Tunisia<br>Madrid, Spain | Yes | No | bacteria | 173±21 |
| 26 | Shiga Toxin-Producing E. Coli (STEC) | Michigan, US<br>Germany<br>UK<br>Kanazawa, Ishikawa, Japan<br>Iceland<br>Netherlands<br>Ireland | Yes | No | bacteria | 225±30 |
| 27 | Shigellosis | Kirkenes, Norway<br>Sichuan, China<br>North Carolina, US<br>Morobe, Papua New Guinea<br>Sweden<br>Virginia, US<br>Honolulu, US<br>Pemagatsel, Bhutan<br>Norway<br>Salzburg, Austria<br>Austria | No | No | bacteria | 179±16 |
| 28 | Tularemia | Germany<br>Utah, US<br>France | Yes | Yes | bacteria | 183±12 |
| 29 | West Nile Virus | Michigan, US<br>Thessaloniki, Greece<br>Romania<br>Italy | Yes | Yes | virus | 145±16 |
| 30 | Yellow Fever | Sudan<br>San Pedro, Paraguay<br>São Paulo, Brazil | Yes | Yes | virus | 92±16 |
| 31 | Zika | San Andrés, Colombia<br>Girardot, Colombia | No | Yes | virus | 120±25 |

**Table S5.**
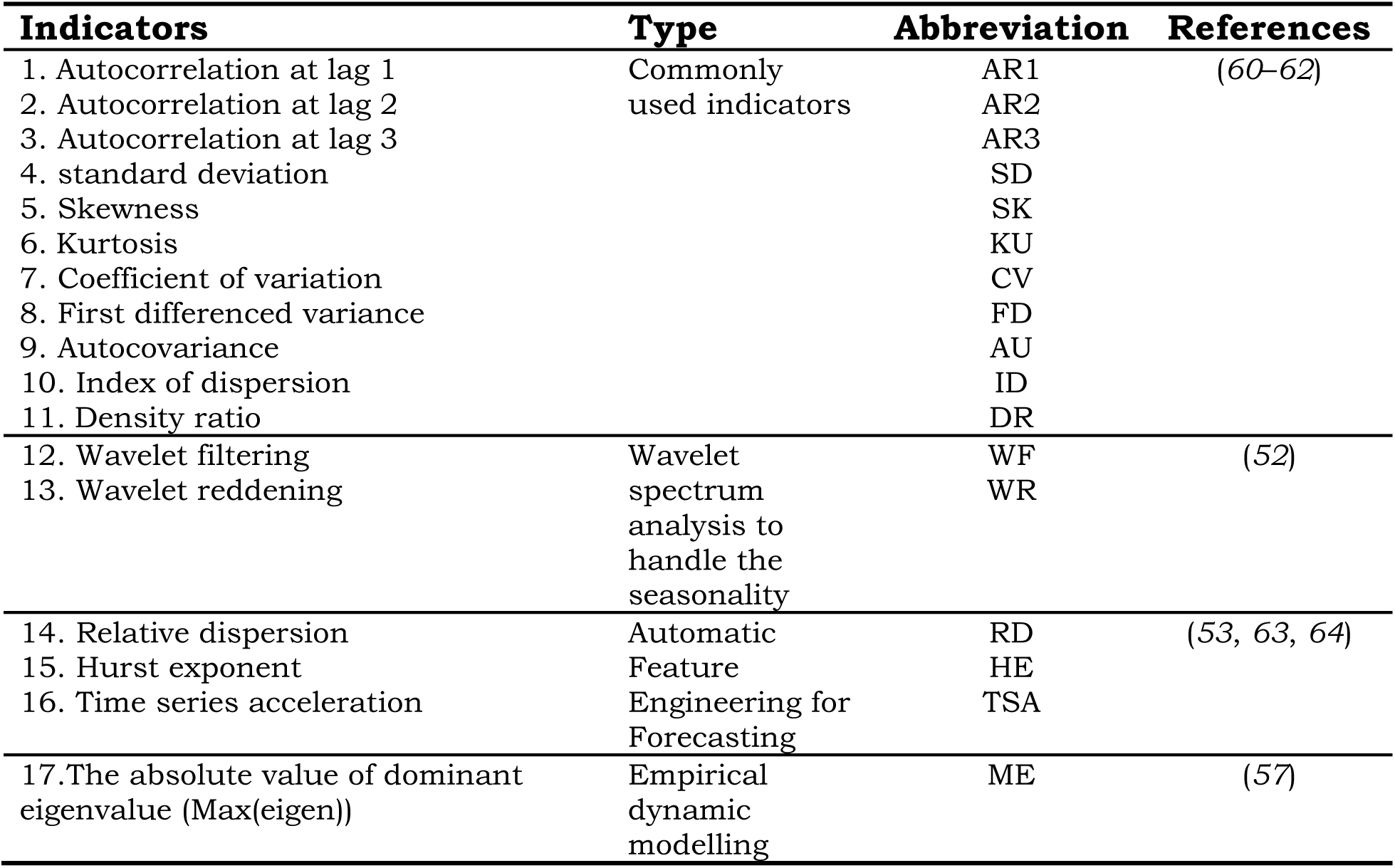
List of 17 resilience indicators (RIs) used to detect critical transitions, and their type, abbreviation, and reference. The details of the 17 resilience indicators RIs see Supplementary information 2.

